# mSWI/SNF complex inhibition sensitizes KRAS-mutant lung cancers to targeted therapies via epithelial-mesenchymal subversion

**DOI:** 10.64898/2026.02.27.708377

**Authors:** Claudia Gentile, William W. Feng, Sean M. Lenahan, Alexander W. Ying, Daren C. Card, Florence T.H. Wu, Nhu-An Pham, Nikolina Radulovich, Pinjiang Mary Cao, Katrina Hueniken, Quan Li, Ming-Sound Tsao, Joseph Kulesza, Madeline M. Hinkley, Lishu Liao, Jeanelle A. Tsai, Jens Köhler, Francesco Facchinetti, Jiaqi Li, Caitlyn Weston, Marie-Anaïs Locquet, Kenneth Ngo, Prafulla C. Gokhale, Adrian G. Sacher, Pasi A. Jänne, Cigall Kadoch

**Affiliations:** Department of Pediatric Oncology, Dana-Farber Cancer Institute and Harvard Medical School, Boston, MA, USA 02215; Broad Institute of MIT and Harvard, Cambridge, MA, USA; Department of Medical Oncology, Dana-Farber Cancer Institute, Boston, MA, USA; Princess Margaret Cancer Centre, University Health Network, Toronto, ON, Canada; Clinician Investigator Program, University of British Columbia, Vancouver, BC, Canada; Department of Medical Biophysics, University of Toronto, ON, Canada; Department of Laboratory Medicine and Pathobiology, University of Toronto, ON, Canada; Belfer Center for Applied Cancer Science, Dana-Farber Cancer Institute, Boston, MA, USA; Department of Immunology, University of Toronto, ON, Canada; Howard Hughes Medical Institute, Chevy Chase, MD, USA

**Author notes:** These authors contributed equally to this work (Claudia Gentile, William W. Feng). Denotes co-corresponding authors ^‡^Correspondence to: Cigall Kadoch, Ph.D. Professor and Meredith and Billy Starr Investigator, Department of Pediatric Oncology Dana-Farber Cancer Institute and Harvard Medical School Institute Member and Epigenomics Program Co-Director, Broad Institute of Harvard and MIT Investigator, Howard Hughes Medical Institute (HHMI) 450 Brookline Avenue, LC-5217 Boston, MA 02215.

## Abstract

Targeted therapies for *KRAS*-mutant non-small lung cancer (NSCLC) have shown promising clinical results, however, incomplete tumoral responses and the inevitable emergence of therapeutic resistance remain critical challenges. Here we identify mSWI/SNF chromatin remodeling complexes as critical determinants of (EMT)-mediated KRAS inhibitor inefficacy and resistance in KRAS G12C lung cancers. Treatment with the clinical-grade SMARCA4/2 inhibitor, FHD-286, dampens EMT-mediated acquired resistance in drug-responsive models and similarly resensitizes drug-refractory models by rewiring mSWI/SNF chromatin localization and activities that modulate epithelial transcriptional programs and cell state. Further, synergistic mSWI/SNF and KRAS inhibitor combination treatment sensitizes non-G12C KRAS-mutant NSCLC cells to pan-RAS and G12D-specific inhibitors. Finally, FHD-286 and sotorasib combination treatment results in potent anti-tumor efficacy in both G12Ci-resistant and - sensitive organoid models and in vivo patient-derived xenograft (PDX) systems. These data nominate mSWI/SNF inhibition as a combination strategy to improve KRAS inhibitor efficacy, response duration, and to mitigate emergence of resistance.

## Introduction

Mutations in *KRAS* are among the most frequent and deleterious genomic alterations across human tumors and occur in one third of all non-small cell lung cancers (NSCLCs)^1^. Mutations in KRAS result in hyperactivation of the mitogen-activated protein kinase (MAPK) and phosphoinositide 3-kinase (PI3K) pathways oncogenic signaling pathways, supporting constitutive activities of downstream protein kinases and effector proteins to promote tumor growth and survival. Decades of focused research in both academic and pharmaceutical sectors have enabled the more recent development of mutant-selective KRAS inhibitors, including the KRAS G12C inhibitors, sotorasib and adagrasib, which are now components of standard-of-care treatment regimens. While early results are promising, long-term durability remains a major challenge as almost all patients exhibit disease relapse within < 8 months^2,3^. These results emphasize the need to identify determinants of resistance and mechanism-based opportunities for therapeutic combination strategies to improve overall efficacy and mitigate the emergence of resistance.

Interestingly, while the acquisition of new genomic alterations including kinase pathway ‘bypass’ mutations often underlie resistance to targeted therapies, nearly half of patients progressing on sotorasib or adagrasib KRAS G12C inhibitors *do not* exhibit a secondary tumor mutation in *KRAS* or other well-established genes linked to bypass survival signaling^4,5^. Given this, chromatin-level (epigenetic) and transcriptional processes have been flagged as dynamic mechanisms that can modulate signaling pathway activities and facilitate the development of therapeutic resistance across a broad spectrum of cancers^6,7^. In addition to KRAS-mutant NSCLC, non-mutational mechanisms of therapeutic resistance are observed in over half of EGFR-mutant NSCLC patients progressing on the third-generation EGFR inhibitor, osimertinib^8,9^. Previous studies by our group have implicated YAP/TEAD-mediated transcriptional rewiring^10^ and mSWI/SNF-mediated chromatin remodeling^11^ as important determinants of osimertinib resistance in EGFR-mutant NSCLC. These data suggest a potentially more broad-spanning use case for inhibiting mSWI/SNF chromatin remodeler activities in the setting of kinase-mutant cancers, especially considering the extensive mutational frequencies of mSWI/SNF components and their roles as top synthetic lethal dependencies in human cancer^12,13^. In addition, small molecules targeting the catalytic engines of these complexes (SMARCA4 (BRG1)/SMARCA2 (BRM) ATPase inhibitors) such as FHD-286 have shown promising results with respect to impact on tumor and tumor microenvironments in Phase I clinical studies to date, making them ideal candidates for consideration in therapeutic combinations^14^.

The context-specific functions of mSWI/SNF complexes in lung cancers of distinct mutational backgrounds remain unclear. *SMARCA4* mutations occur in approximately 11% of NSCLCs, in which SMARCA2, the SMARCA4 subunit paralog, is the top-ranked synthetic lethal vulnerability^15–17^. Notably, *SMARCA4* mutations rarely co-occur with *EGFR* mutations or *ALK* rearrangements, but more frequently co-occur with *KRAS* mutations, implicating an interplay-either via separate or cooperative pathways-between hyperactive oncogenic kinases and chromatin remodeling effectors ^18,19^. Patients harboring tumors with dual *SMARCA4* and *KRAS* mutations demonstrate poorer responses to KRAS-directed therapies and worse overall survival relative to patients with tumors lacking *SMARCA4* mutations^20^. Genetically engineered mouse models (GEMMs) have shed light on the seemingly paradoxical role of SMARCA4 in tumor suppression; SMARCA4 depletion inhibits lung tumor progression but its deficiency can also give rise to advanced lung tumors and metastasis, depending on the cell type in which loss of SMARCA4 function occurs^19^. The divergent pro- and anti-tumorigenic activities of BAF complexes in lung adenocarcinoma (LUAD) initiation and progression^19^ have similarly been reported in pancreatic GEMMs in which *Smarca4* knockout prevents initiation of pancreatic intraepithelial neoplasia yet promotes oncogenic transformation in pancreatic duct cells^21^. Together, these results demonstrate the highly context-dependent activities of mSWI/SNF complexes as both tumor suppressors and oncogenes, and motivating a need to better define their roles in KRAS-mutant NSCLC.

Here we reveal the functions of mSWI/SNF complexes in mediating epithelial-mesenchymal transition (EMT) transcriptional programs in KRAS-mutant lung cancer. We identify mSWI/SNF-directed control of the receptor tyrosine kinase, AXL, as a key mediator of resistance to KRAS inhibitor treatment via promotion of EMT which is opposed by mSWI/SNF inhibition. Pharmacologic SMARCA4/2 ATPase inhibition alone ablates chromatin accessibility and expression of EMT-associated gene loci including *AXL*, rewiring cells toward an epithelial cell state and enhances the activity of KRAS inhibitors in a subset of KRAS-mutant cells. We find that this mechanism is intact across diverse models, with mSWI/SNF inhibition sensitizing various KRAS-mutant NSCLCs to the pan-RAS inhibitor, RMC-6236, through analogous means as well as resensitizing resistant cells to KRAS-directed therapies. Importantly, combinatorial inhibition of mSWI/SNF and KRAS elicits enhanced anti-tumor efficacy relative to either agent alone in xenograft-derived organoid (XDO) and patient-derived xenograft (PDX) models. Our findings position mSWI/SNF-mediated maintenance of EMT as a pharmacologically targetable therapeutic vulnerability in a spectrum of KRAS-mutant lung cancers, providing important rationale for clinical evaluation of combination mSWI/SNF and KRAS inhibition.

## Results

### Inhibition of mSWI/SNF chromatin remodeling complexes synergizes with sotorasib and adagrasib in KRAS-mutant lung cancer cells

Upstream transcriptional regulator analyses using RNA-seq data from n=142 primary human KRAS-mutant lung adenocarcinomas (LUAD) revealed several transcription factors (TFs) and epigenetic proteins as top predicted regulators (**Figure 1A**). Genes encoding canonical BAF (cBAF) complex subunits ARID1A and SMARCA4 were ranked 7th and 10th, respectively, among all transcriptional regulators in KRAS-mutant primary tumors (**Figures 1A, S1A-B**). Similar results were obtained examining gene expression in human KRAS-mutant cell lines from the DepMap CCLE database (**Figure S1C**), with mSWI/SNF genes again identified as top transcriptional regulators (**Figure S1D**), suggesting mSWI/SNF complexes may play key gene regulatory roles in KRAS-mutant lung cancer.

**Figure 1.**
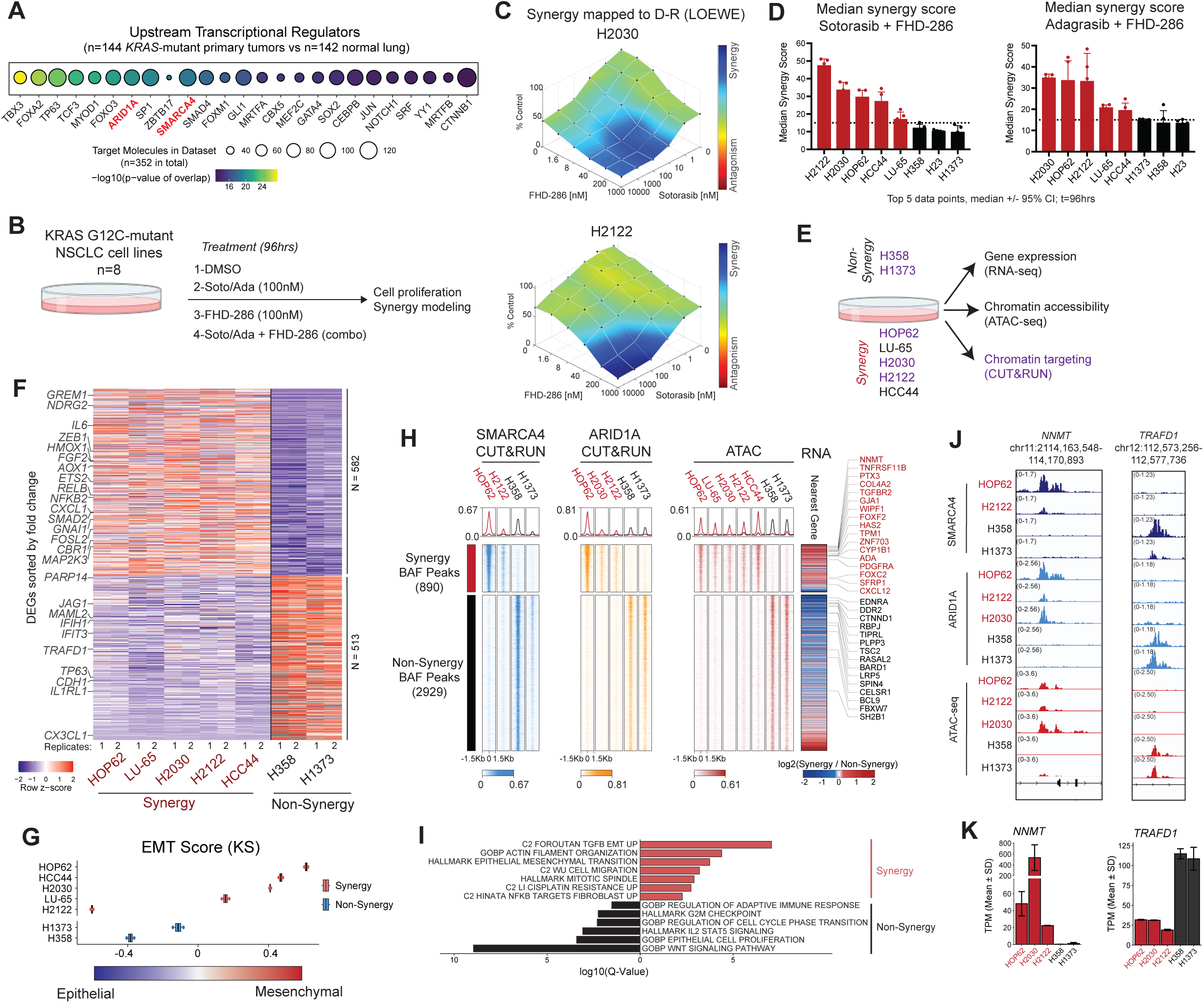
mSWI/SNF complex inhibition synergizes with sotorasib to govern KRAS-mutant cell gene expression signatures. A. Upstream transcriptional regulator analysis (IPA) performed on KRAS-mutant NSCLC primary tumors (TCGA; n=142) versus normal lung epithelium samples (GTEx, n=144) depicting top 25 factors. B. Schematic for evaluating single-agent and combination treatment using KRAS G12C inhibitors, sotorasib (Soto) and adagrasib (Ada), and the SMARCA4/2 inhibitor, FHD-286. C. Combenefit (LOEWE) modeling analysis of synergy assays performed with sotorasib and FHD-286 in H2030 and H2122 KRAS G12C cell lines. D. Synergy scores (LOEWE modeling) for n=8 cell lines treated with sotorasib or adagrasib + FHD-286 reveals ‘synergy’ (red) and ‘non-synergy’ (black) cell lines at t= 96 hours. n=5 experimental replicates; median plotted -/+ 95% CI; dotted line represents synergy cutoff >/= 15; see also Figure S1H-I). E. Schematic for genomics-centered experiments in KRAS G12C ‘synergy’ and ‘non-synergy’ cell lines. Purple highlighted cell lines are selected for CUT&RUN. F. Heatmap depicting up- and down-regulated genes in synergy and non-synergy lines; selected genes are labeled. G. Epithelial to mesenchymal transition (EMT) score for KRAS-mutant cell lines using the KS (Kolmogorov-Smirnov) method. H. Heatmaps of synergy and non-synergy specific mSWI/SNF occupancy measured by CUT&RUN for SMARCA4 and ARID1A and corresponding chromatin accessibility (ATAC). Nearest gene expression (RNA-seq) is represented as fold change between synergy and non-synergy cell lines; EMT related genes are labeled in red and G2M/WNT signaling non-synergy genes are labeled in black. I. Overrepresentation gene set enrichment analysis using curated gene pathways from MSigDB for complex-occupied synergy-specific and non-synergy specific genes from (H). J. Representative tracks at the NNMT and TRAFD1 gene loci showing mSWI/SNF complex occupancy (ARID1A and SMARCA4 CUT&RUN) and ATAC-seq. K. Gene expression (TPM) bar plots of NNMT and TRAFD1 gene expression across cell lines.

We next sought to evaluate the impact of combined mSWI/SNF and KRAS inhibition on KRAS-mutant cell lines. Selecting n=8 KRAS G12C-mutant cell lines with various co-mutational backgrounds representative of the clinical spectrum of disease (**Figure S1E**), we defined IC_50_ values for KRAS inhibition using either sotorasib or adagrasib, and for mSWI/SNF complex inhibition using the clinical-grade dual SMARCA4/A2 ATPase inhibitor, FHD-286 (**Figure S1F**). Sotorasib and adagrasib monotherapy exhibited relatively low potency, reflective of their sub-optimal activity in patients; FHD-286 demonstrated poor single agent efficacy, in agreement with their lack of overt mSWI/SNF fitness dependencies (**Figure S1F**). Indeed, comprehensive cell line profiling of FHD-286 (using PRISM ^22,23^), revealed that only 7 of 24 KRAS-mutant NSCLC lines show meaningful inhibition of proliferation (AUC > 0.8) upon FHD-286 treatment (**Figure S1G and Table S1**).

As FHD-286 treatment alone did not have a major proliferative impact, we evaluated drug synergy of FHD-286 plus KRAS G12C inhibitors using Combenefit ^24^ (**Figure 1B**). Notably, combination treatment generated substantial synergy in 5 of 8 total lines evaluated (**Figures 1C-D, S1H-I**). No drug synergy was observed in the H23 cell line (**Figures S1H-I**), which we confirmed to be *SMARCA2* and *SMARCA4* dual-deficient by Whole Exome Sequencing (WES) (**Table S2**). Notably, H2030 displayed a striking degree of synergy despite its purported bi-allelic deletion of *SMARCA4*, which we validated by WES (**Figures 1C and Table S2**), suggesting FHD-286 targeting of the remaining SMARCA2 ATPase paralog. Indeed, shRNA-mediated knockdown of SMARCA2 in combination with sotorasib in H2030 cells resulted in similar attenuation of cell proliferation, further validating the observed synergy (**Figures S1J-K**). Together, these data show that while FHD-286-mediated ATPase inhibition has minimal impact as a monotherapy in KRAS G12C-mutant lung cancers, combination treatment of KRAS G12C inhibitors with FHD-286 shows a synergistic anti-proliferative impact in a subset of KRAS G12C NSCLC cell lines.

### mSWI/SNF complexes govern an EMT signature specific to the synergy response in a subset of KRAS-mutant cell lines

We next sought to define potential mSWI/SNF-mediated chromatin-level features that distinguish KRAS-mutant lines exhibiting “synergy” or “non-synergy” with combination FHD-286 and KRAS inhibition. We first performed RNA-seq across the panel of KRAS-mutant cell lines, apart from H23 owing to dual SMARCA4/A2 loss (**Figure 1E**). Notably, we identified a collection of differentially expressed genes (DEGs) that were up- and down-regulated specifically in either synergy or non-synergy cell lines (**Figure 1F**). Gene set enrichment analysis (GSEA) revealed strong positive enrichment for epithelial-to-mesenchymal transition (EMT) and negative enrichment for the interferon alpha response gene signatures in synergy cell lines compared to non-synergy cell lines (**Figure S1L**). As examples, we found key EMT genes preferentially expressed in synergy cell lines, including *IGF1R*, *TWIST1*, and *ZEB1,* whereas IFN-α genes such as *CX3CL1*, *IFIH1*, and *MAML2* were differentially upregulated in non-synergy cell lines (**Figure S1M**). The EMT status of each cell line was mapped across an EMT spectrum as defined using the K-S method from a previously described gene set signature ^25,26^ (**Figure 1G**). Cell lines exhibiting synergy with FHD-286 nearly uniformly scored as mesenchymal, except for H2122 cells, potentially attributed to their enrichment for an adenocarcinoma-to-squamous cell carcinoma gene signature ^27^, another lineage plastic state responsive to mSWI/SNF inhibition (**Figure S1N**).

Interestingly, we noted that 4 of 5 cell lines demonstrating drug synergy (H2122, H2030, HCC44 and LU-65) are classified as the *STK11*/*LKB1*-mutant (“KL”) subtype. We therefore sought to determine if synergy response could be mechanistically explained by *STK11* mutational status. However, neither CRISPR-mediated KO of *STK11* in “KP” lines (*p53*-mutant, *STK11/LKB1*-wildtype), H358 and H1373, nor rescue of *STK11* expression in “KL” lines (H2122, HCC44, and H2030) resulted in meaningful changes to combination FHD-286 and sotorasib synergy response, despite achieving expected impacts on downstream phosphorylation status of ACC and RAPTOR (**Figure S1O-P**). These results point to predictive transcriptional signatures, rather than mutational subtype, as important determinants of KRAS inhibitor and mSWI/SNF inhibitor synergy. Further, the one synergy cell line, HOP62 to be considered a “KP” subtype exhibited a strong EMT signature, likely underlying its response to combination treatment (**Figures 1G, S1E, S1L**).

We next profiled the genomic occupancy of mSWI/SNF complexes in a selected set of synergy and non-synergy cell lines using SMARCA4, ARID1A and SMARCC1 CUT&RUN (**Figure 1E, S1S**). In parallel, we performed ATAC-seq in each cell line to define genome-wide chromatin accessibility (**Figure 1E**). As expected, mSWI/SNF occupancy strongly correlated with chromatin accessibility (ATAC-seq signal) genome-wide, as well as with H3K27ac occupancy (**Figures S1S-T**). Importantly, we identified specialized sites of SMARCA4 and ARID1A occupancy across all synergy lines relative to non-synergy lines which mapped to sites of enhanced chromatin accessibility (ATAC-seq), including loci containing genes uniquely activated in synergy versus non-synergy lines (**Figures 1H, S1U**). As examples, synergy-specific accessible mSWI/SNF target sites enriched for gene sets controlling EMT, cytoskeletal organization and cell migration, among others, as exemplified at the *NNMT* locus (**Figures 1I-K**). In contrast, the non-synergy-specific mSWI/SNF-occupied sites enriched for genes involved in WNT signaling and cell cycle regulation, exemplified by the *TRAFD1*, *SELL* and *TP63* loci (**Figures 1I-K, S1U**). Together, these data suggest a unique set of features controlled by mSWI/SNF targeting and activity that govern FHD-286 synergy and non-synergy responses in KRAS-mutant cell lines.

### Combination sotorasib and FHD-286 treatments attenuates EMT and cell cycle progression in KRAS G12C-mutant cells

Given our results implicating mSWI/SNF complexes in the regulation of synergy- and non-synergy-specific chromatin features, we next sought to define their contributions in supporting KRAS signaling. We treated two synergy cell lines (H2122 and H2030) and two non-synergy cell lines (H358 and H1373) with sotorasib, FHD-286, or the combination of both drugs and assessed impact on gene expression and chromatin accessibility (**Figure 2A**). Treatment with FHD-286 alone resulted in minimal impacts on differential gene expression (relative to sotorasib or combo) in all cell lines, while combination treatment resulted in greater numbers of differentially down- and up- regulated genes, largely associated with loci occupied by mSWI/SNF complexes (**Figures S2A-C**). Overall, combination treatment resulted in unique sets of differentially expressed genes (compared to sotorasib or FHD-286 alone) (**Figures 2B, S2D**). Importantly, in H2030 and H2122 synergy cell lines, C1 gene clusters (combo upregulated) were enriched for genes involved in transcriptional regulation (eg. *SP1*), while genes in cluster 2 (combo downregulated) enriched for epithelial cell migration (e.g. *PTK2*) and EMT (e.g. FOS), in each respective synergy cell line (**Figure 2B**). Moreover, a subset of differentially expressed genes demonstrated an enhanced downregulation upon combination treatment as compared to sotorasib treatment alone (C3 genes) across all cell lines (**Figure 2B, S2D**), demonstrating that mSWI/SNF inhibition increases the depth of response observe with KRAS inhibition (sotorasib). In synergy cell lines, these gene sets enriched for epithelial cell differentiation and TGFB1 response (e.g. *NOTCH1*, *AKT1*, *NF2*) (**Figure 2B**). In contrast, in non-synergy cell lines, combination treatment enhanced the downregulation of genes involved in DNA damage checkpoints as exemplified by *TOP3A* and *UIMC1* (**Figure S2D**).

**Figure 2.**
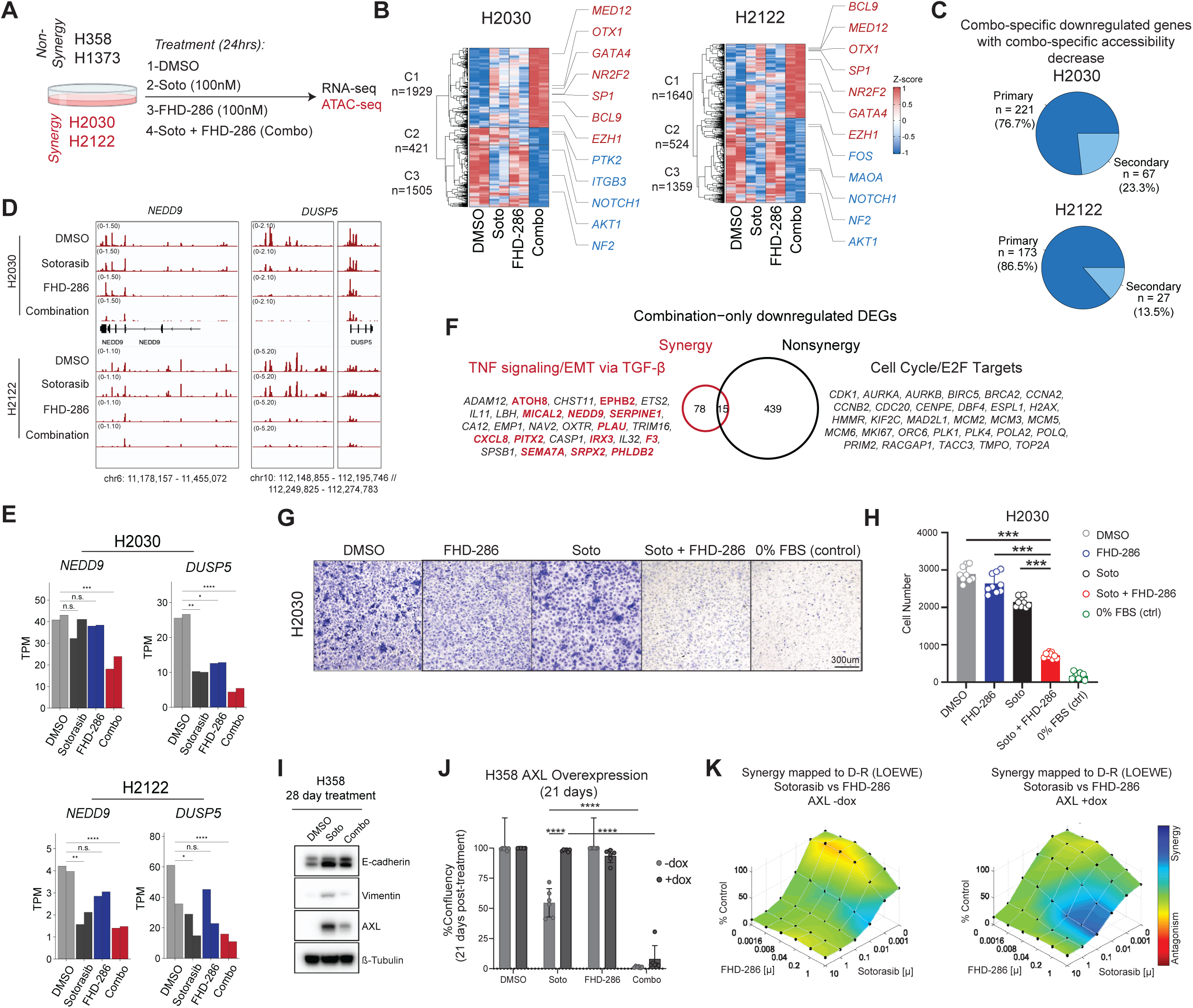
FHD-286-mediated subversion of epithelial-mesenchymal transition potentiates combination synergy in KRAS-mutant NSCLC cells. A.Schematic for genomics-centered experiments following drug treatment of non-synergy and synergy cell lines. ATAC-seq was performed only in synergy lines. B. Row z-score normalized heatmap of gene expression following sotorasib, FHD-286 and combination treatments in H2030 and H2122 cells. Genes displayed represent the top ten percent of combination-specific up- or down-regulated genes. Gene examples are labeled. C. Pie charts representing percentage of combination specific down-regulated genes with concordant chromatin accessibility (ATAC-seq) changes upon combination treatment (primary targets) in H2030 and H2122. Nearest ATAC-seq peak was assessed within a 30kb window of TSS. D. IGV snapshots of ATAC-seq tracks in H2030 and H2122 cell lines treated with DMSO, sotorasib, FHD-286 and combination over the NEDD9 and DUSP5 loci. E. Gene expression (TPM) bar graphs for NEDD9 and DUSP5 across indicated treatment conditions. Significance is displayed using DESeq2 adjusted p-value. F. Venn diagram overlap of combination only down-regulated genes in synergy and non-synergy cell lines (resulting from subtracting sotorasib downregulated genes). Relevant genes enriched for TNF signaling/EMT via TFG-B and for Cell Cycle/E2F pathways are listed. Genes involed in regulation of migration are labeled in red. G. Transwell migration assay of H2030 cells pre-treated with DMSO, 100nM soto, 100nM FHD-286 or combo for 7 days before seeding into inserts for assay. Representative images taken 72hrs after seeding (10X magnification) are shown from a single experiment from n=3 independent experiments. H. Bar graph of number of cells that migrated through Transwells from 3 distinct fields of view from biological replicates. Statistical significance is assessed by one-way ANOVA with Sidak’s multiple comparisons test. *** p < 0.0001. I. Immunoblot of EMT markers E-Cadherin, vimentin, and AXL in H358 cells following 28 day treatment of sotorasib or combo. B-tubulin is the loading control. J. Bar charts of % confluency of long-term viability assay for H358 cells over-expressing AXL (14 days) followed by sotorasib, FHD-286, or combination drug treatment over 21 days (with continuous +/-dox treatment). Significance is denoted by asterisks and determined using two-way Anova with Tukey’s multiple comparisons. K. Combenefit synergy assays performed with sotorasib and FHD-286 in H358 cells treated without or with dox to induce AXL overexpression. Dox treatment was done for 14 days followed by 96hrs of drug treatment.

Further, genes downregulated following combination treatment were associated with combination-specific decreases in chromatin accessibility (primary targets) at over 75% of downregulated targets in synergy cell lines H2122 and H2030, as exemplified at the *NEDD9*, and *DUSP5* loci (**Figures 2C-E, S2E**). Gains in chromatin accessibility were also detected to a lesser degree and corresponded to upregulated genes (primary targets), suggesting a reshuffling of active chromatin architecture upon combined inhibition of mSWI/SNF ATPase and KRAS activities (**Figures S2E-F**).

Next, subtracting the intersect of up- or down-regulated genes across synergy and non-synergy cell lines following sotorasib treatment from those deregulated in combination treatment, revealed combo-only impacts (**Figures 2F, S2G**). Combo-only down-regulated genes in synergy cell lines enriched predominantly for TNF signaling and EMT via TGF-β signaling and at the protein level for classic EMT markers (**Figure 2F and S2H**). Conversely, combo-only down-regulated genes in non-synergy cell lines enriched for cell cyle pathways and E2F targets (**Figure 2F**). These results are consistent with mSWI/SNF-specific occupancy at targets which distinguish synergy and non-synergy cell lines at the transcriptional and chromatin levels (**Figure 1H**). Further, up-regulated combo-only gene targets were distinct between synergy and non-synergy cell lines, but enriched for various pathways and processes (**Figure S2G**). Together these findings support a model in which concomitant shut down of mSWI/SNF ATP-dependent chromatin remodeling and mutant KRAS signaling are required to rewire gene expression and chromatin states that potentiate KRAS inhibitor response.

Within synergy-specific combo-only downregulated genes, we identified a large subset of genes enriching for regulation of cell migration (**Figure 2F**), a well-established hallmark of cells undergoing EMT ^28^. Notably, transwell migration assays on mesenchymal H2030 cells revealed significant decreases in the migratory potential of H2030 cells following 7 days combination FHD-286 and sotorasib treatment, an effect not achieved by either single agent alone (**Figures 2G-H**). Together, these data suggest that mSWI/SNF inhibition antagonizes mesenchymal cellular programming.

In parallel, to evaluate cell cycle, we employed a genetically encoded Fluorescent Ubiquitination-based Cell Cycle Indicator (FUCCI) reporter system ^29^ and performed time course cell cycle analysis following KRAS and BAF inhibitor treatment in H2030 (synergy) cells and H1373 (non-synergy) cells. In H2030 cells, we observed that while sotorasib alone elicits robust G1 arrest within 1 day of treatment, this was reversed within 7 days, whereas cells treated with combo exhibited a durable increase in %G1 cells and a concomitant reduction in %S and %G2M populations (**Figure S2I**), supporting the synergistic responses observed. Further, while FHD-286 alone had minimal impact on cell cycle progression of H2030 cells, it was as effective as sotorasib in attenuating cell cycle progression in H1373 (**Figure S2I**), leading to negligible benefit when used in combination. These results provide evidence that the absence of synergy in H1373 is likely due to redundant effects of single agent and combinatorial KRAS and BAF inhibition, while in H2030 “synergy” cells, combination inhibitor treatment elicited a clear benefit over either agent alone.

To validate the role for mSWI/SNF complexes in driving EMT to facilitate a synergistic response to combination treatment, we tested whether mSWI/SNF complex activity is sufficient for the EMT state change. As H358 cells exhibit a more epithelial-skewed cellular state (**Figure 1G**), it represents an optimal setting to interrogate the ability of the BAF complexes to drive EMT. Strikingly, following treatment of H358 cells with sotorasib for 28 days, expression of canonical EMT mesenchymal markers AXL and vimentin markedly increased, whereas co-treatment with FHD-286 largely attenuated this effect (**Figure 2I**). Similarly, in H2030 cells bearing an EMT-high transcriptional signature, FHD-286 in combination with sotorasib resulted in reduced chromatin accessibility at the *AXL* locus (**Figure S2J**). As AXL is a well-known driver of EMT-mediated resistance in lung cancer ^30–32^ we next mimicked EMT-mediated therapeutic resistance by overexpressing AXL in H358 cells (**Figure S2K**). While overexpression of AXL alone had no effect on growth rate as compared to the no dox DMSO control, AXL overexpression conferred a substantial proliferative advantage in sotorasib-treated cells over 21 days (**Figure 2J, S2L**). Notably, this sotorasib-refractory effect was durably rescued by FHD-286 over 21 days of treatment (**Figure 2J, S2L**) and was confirmed to be synergistic (**Figure 2K)**. These results demonstrate that elevated AXL expression confers decreased sensitivity to sotorasib and that FHD-286 promotes re-sensitization to sotorasib, in part through opposing an AXL-induced EMT-axis.

### Inhibition of mSWI/SNF complexes displays broad-spectrum efficacy in diverse KRAS-mutant lung cancers

We next performed longitudinal live cell imaging assays to explore the impact of combination inhibition of mSWI/SNF and KRAS on the durability of drug response. Nearly all drug-naïve KRAS G12C cellular models exhibited rapid relapses on sotorasib single-agent treatment, except for H358 (**Figure 3A**), mirroring short response durability in patients. While FHD-286 treatment alone only modestly attenuated proliferation across cell lines, the combination of sotorasib and FHD-286 elicited a striking attenuation of cellular outgrowth in H2122 and H2030 cell lines (**Figure 3A**), concordant with marked synergy observed at 96 hours (**Figure 1C**). The superiority of the drug combination over either single agent was not limited to “synergy” cell lines; indeed, added benefit of FHD-286 manifested over time in H358 and H1373 cell lines as well, despite the absence of drug synergy in short-term assays (i.e. non-synergy cell lines) (**Figures 3A, 1D, S1H-I).** To note, whereas phospho-ERK levels quickly rebounds within days following sotorasib treatment across cell lines, combination treatment with FHD-286 results in pERK suppression to degrees achieved by KRAS and MEK inhibitor combinations (**Figure S3A**). These results demonstrate that pharmacologic inhibition of mSWI/SNF complexes potentiates KRAS inhibitor treatment efficacy significantly beyond what is achieved with single-agent treatment.

**Figure 3.**
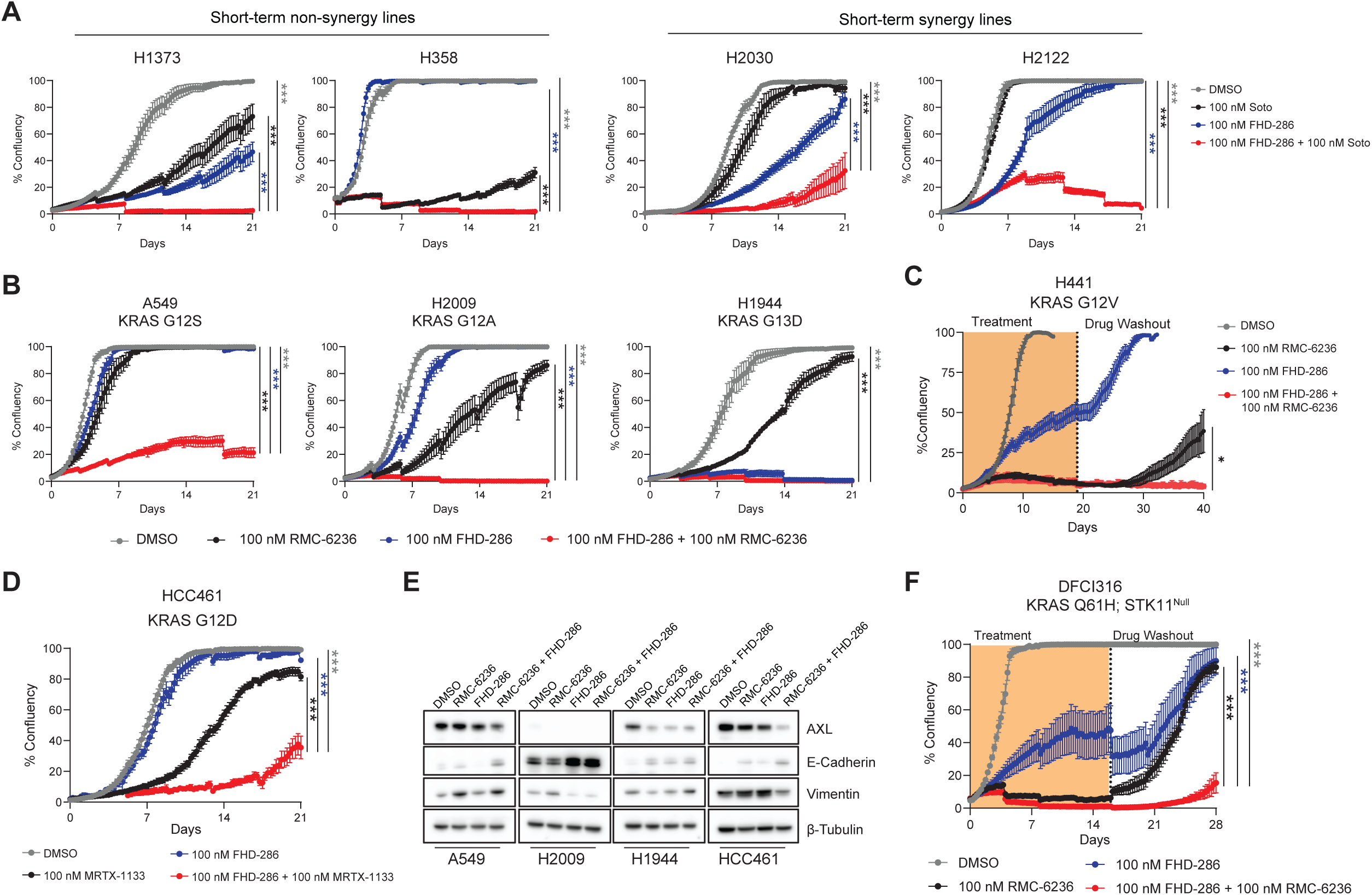
mSWI/SNF inhibition broadly potentiates activity of pan-RAS and KRAS G12D-specific inhibitors. A. Confluence measurements of H1373 and H358 (non-synergy) and H2030 and H2122 (synergy) cells with indicated treatment conditions were measured once every 6 hours over 21 days, media replenished every 4 days. B-C. Confluence of cells exposed to indicated treatments was measured once every 6 hours over 21 days for A549, H2009, and H1944. (C) Drugs were washed out from H441 and cells were continuously imaged until day 40 to evaluate for signs of cellular regrowth. Media was replenished every 4 days. D. Confluence of HCC461 cells exposed to the indicated treatments were measured once every 6 hours for 21 days. E. Western blot analysis for EMT markers after 96 hours of treatment with DMSO, 100 nM sotorasib, 100 nM FHD-286, or 100 nM sotorasib + 100 nM FHD-286. F. Confluence of DFCI316 cells exposed to the indicated treatments was measured once every 6 hours over 28 days. Drugs were washed out from on day 16 and cells were continuously imaged to evaluate for signs of cellular regrowth. Media was replenished every 4 days. One-way ANOVA was used to compare groups at endpoint for (A-D), and (F). ***p < 0.0005, *p < 0.05. Data represent mean +/- SEM of n=4 replicates.

While the G12C mutation is the most frequent KRAS alteration occurring in LUAD tumors (**Figure S1A**), we next sought to evaluate whether these results could be extended to KRAS-mutant lung cancer models harboring non-G12C KRAS mutations with varying co-mutational backgrounds and EMT status (**Figures S3B-C**). Excitingly, we found that FHD-286 strongly sensitizes A549 (KRAS G12S) and H2009 (KRAS G12A) cell lines to the clinical-grade pan-RAS inhibitor, RMC-6236, and exhibits marked single agent activity in H1944 (KRAS G13D) (**Figure 3B**). Further, despite H441 (KRAS G12V) cells exhibiting robust sensitivity to RMC-6236 alone, following drug discontinuation, H441 cells quickly relapsed (**Figure 3C**). However, combination treatment with FHD-286 led to complete and sustained responses (**Figure 3C**). Notably, FHD-286 also enhanced durability of response for the KRAS G12D-specific inhibitor MRTX-1133^33^ in HCC461 (KRAS G12D) cells (**Figure 3D**). Across these cell lines, FHD-286 combination treatments decreased AXL expression and/or enhanced E-cadherin expression, mirroring our observations in the KRAS G12C setting (**Figure 3E**). In agreement, long-term treatment (28 days) of H441 cells with RMC-6236 induced AXL expression, an effect largely dampened upon combination treatment with FHD-286(**Figure S3D**). Further, treatment of A549, H2009, and H1944 with the selective AXL inhibitor, bemcentinib ^34^, recapitulated trends seen with FHD-286 treatment conditions (**Figure S3E**). These data provide further evidence for mSWI/SNF-mediated regulation of AXL (and more broadly, the EMT axis) as a critical determinant of treatment failure to RAS inhibitors across a spectrum of distinct mutational contexts and position mSWI/SNF inhibition as a favorable strategy target AXL, given the current lack of effective AXL inhibitors in the clinic (NCT05469178,^35,36^).

Lastly, we evaluated the impact of FHD-286 treatment in a patient-derived cell line model harboring a KRAS^Q61H^ mutation established in our laboratory, DFCI316 ^37^. Although DFCI316 exhibits strong sensitivity to RMC-6236, co-treatment with FHD-286 greatly enhanced the durability and degree of response as evidenced by significantly delayed regrowth kinetics following drug washout (**Figure 3F**). Taken together, these data extend the pivotal role of mSWI/SNF inhibition in enhancing treatment durability, in part via an EMT axis, to an expanded cohort of KRAS-driven lung cancers treated with pan-RAS inhibitors.

### FHD-286 resensitizes drug-refractory tumor-derived organoid models to KRAS inhibition

As FHD-286-mediated mSWI/SNF inhibition increased the durability of response to KRAS inhibitors over time, we next evaluated its role in modulating drug sensitivity in models of stable resistance to KRAS-targeted therapies. To this end, we derived sotorasib- and adagrasib-refractory derivatives of H358 cells following 3 months of culture in the presence of either drug (**Figure 4A**), confirmed by marked shifts in IC_50_ upon KRAS inhibitor treatment (**Figure 4B**). H358 sotorasib-resistant cells (H358SR) demonstrated robust induction of AXL and vimentin which was completely subverted upon FHD-286 treatment (**Figure 4C**). We thus performed SMARCA4 and SMARCC1 CUT&RUN to define mSWI/SNF targeting in sotorasib-resistant H358 cells and to evaluate whether an EMT signature is found in the resistant state of these cells. Notably, we found that mSWI/SNF targeting in H358 resistant cells differs substantially from the retargeting observed following acute (24hr) sotorasib treatment (**Figure 4D**). Interestingly, these distinct targeting patterns were associated with specific gene pathways (**Figure 4E**), revealing that mSWI/SNF retargeting is a distinguishing feature occurring upon the acquisition of sotorasib resistance. While complex occupancy upon acute sotorasib treatment is associated with gene pathways such as cytoskeletal organization, cell junction assembly and epithelial cell differentiation, in the resistant state, mSWI/SNF occupancy is enriched at gene loci associated with cell motility, EMT, TNFA signaling via NFKB, and integrin signaling, as evidenced at the *ITGB3* and *COL6A1* loci (**Figure 4E-F, S4A**). These data demonstrate that the unique retargeting of mSWI/SNF complexes in resistant cells is congruent with mSWI/SNF-specific occupancy in ‘synergy’ cell lines, and supports pathways upholding the EMT-mediated drug resistance.

**Figure 4.**
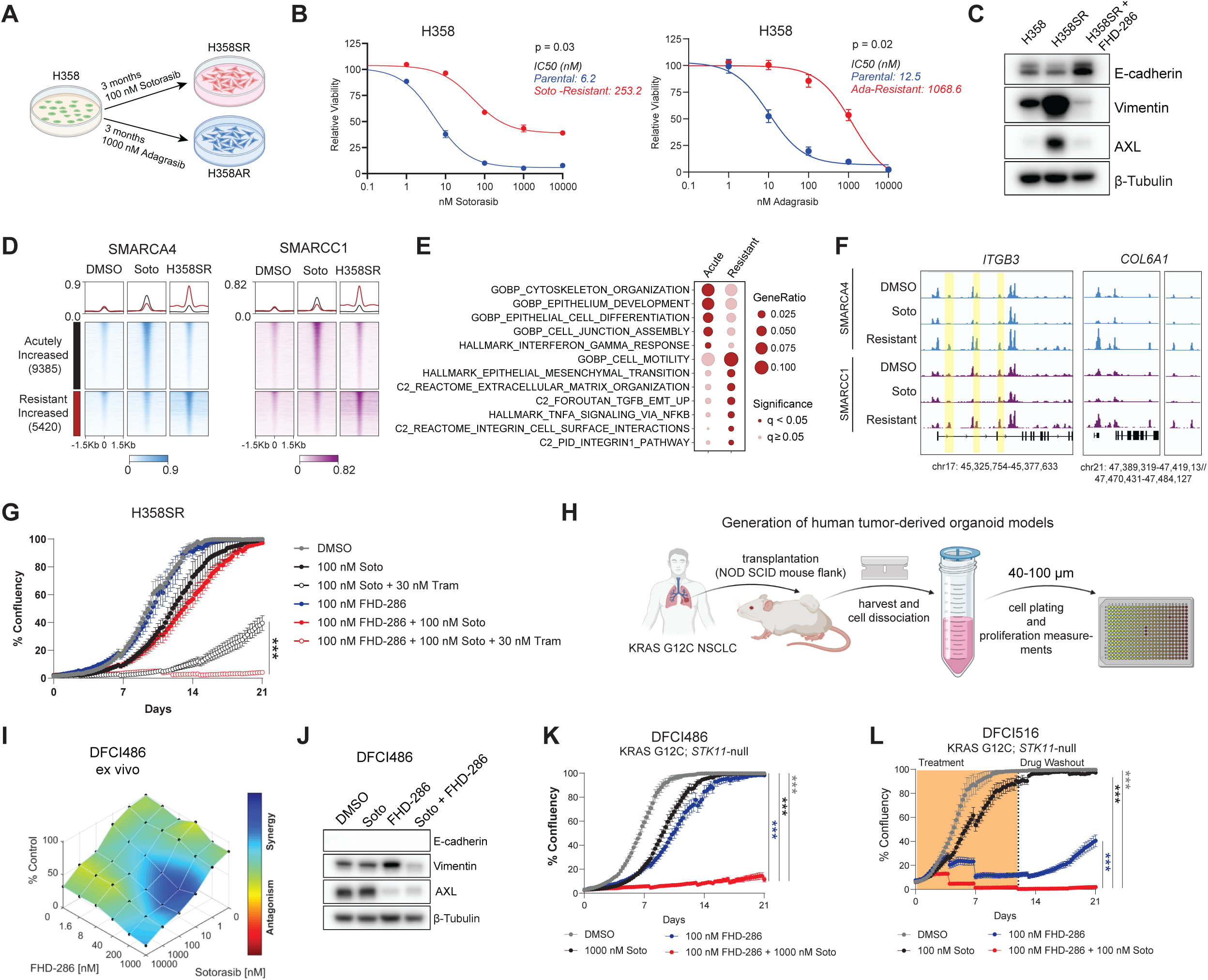
mSWI/SNF inhibition resensitizes drug-refractory lung cancer models to KRAS inhibition. A.Schematic of how sotorasib- and adagrasib-resistant cells were generated. B. IC50 curves comparing sensitivity of parental H358 cells to sotorasib (left) and adagrasib (right). Data represent mean +/- SD from N=3 independent experiments. Welch’s t-test was used to compare IC50s between experiments. C. Western blot comparing expression of EMT markers in H358, H358SR (in presence of 100 nm sotorasib), and H358SR (in presence of 100 nM sotorasib) treated with 100 nM FHD-286 for 96 hrs. D. Heatmaps of SMARCA4 and SMARCC1 CUT&RUN signal across DMSO, 24hr sotorasib treatment (soto) and sotorasib resistant (H358SR) conditions in H358 cells. E. Pathway analysis of genes within 30kB of mSWI/SNF-gained sites in acute soto and resistant conditions. F. IGV track examples of SMARCA4 and SMARCC1 occupancy (CUT&RUN) at ITGB3 and COL6A1 gene loci in H358 cells treated with DMSO, 100nM soto and in H358SR cells. Increased peaks of interest are highlighted. G. Following 1 week sotorasib washout, H358SR cells challenged with the indicated treatments. Wells were imaged once every 6 hours over 21 days. Media was replenished every 4 days. H. Schematic of ex vivo tumor spheroid derivation. See methods for additional details. I. DFCI486 ex vivo tumor spheroids were challenged with drug combination matrices with dose titrations of sotorasib and FHD-286 in ULA 384-well plates. Following 96 hours, 3D Cell Titer Glo (CTG) assay was performed and drug synergy was calculated using Combenefit software. Processed tumor material 40-100 µm was frozen and used for experimental repetitions. One representative experiment out of three is shown. J. Western blot evaluating EMT markers in DFCI486 patient-derived cell line after 96 hours. E-cadherin not detected. K. Confluence of DFCI486 cells treated with the indicated treatments was measured once every 6 hours over 21 days. Media was replenished every 4 days. L. Confluence of DFIC516 cells treated with the indicated treatments was measured once every 6 hours over 21 days. Drug treatment was discontinued (drug washout) on day 12 and wells were continuously scanned for signs of cellular regrowth. Media was replenished every 4 days. One-way ANOVA was used to compare groups at endpoint for (G), and (K-LL). ***p < 0.0005. Data represent mean +/- SEM of n=4 replicates.

Next, while FHD-286 treatment was able to efficiently redirect cells towards an E-cadherin-high, epithelial-leaning cell state (**Figure 4C**), H358SR cells were only durably resensitized to sotorasib upon co-treatment with FHD-286 and the MEK inhibitor, trametinib (**Figure 4G**). Interestingly, H358 adagrasib-resistant cells (H358AR) did not undergo induction of AXL or vimentin expression but FHD-286 similarly guided cells to an E-cadherin enriched state and durably sensitized cells to adagrasib and trametinib (**Figure S4B-C**). Our results recapitulate the central role of pERK reactivation as an adaptive bypass mechanism to KRAS inhibitors seen by others ^38,39^ and position mSWI/SNF ATPase inhibition as a novel vertical inhibition node to be leveraged to enhance the durability of KRAS-directed therapies akin to the KRAS/MEK combinations currently being tested in ongoing human trials ^40^.

We next leveraged *ex vivo* tumor spheroid short-term cultures ^41,42^ from two patient-derived xenograft models harboring KRAS G12C alterations, DFCI486 and DFCI491 (**Figure 4H)**, enabling us to perform high-throughput screening with a range of drug concentrations otherwise unfeasible to pragmatically assay at scale in mice. Both DFCI486 and DFCI491 *ex vivo* tumor spheroid models demonstrated robust drug synergy between FHD-286 and sotorasib, associated with decreases in protein levels of AXL and Vimentin in DFCI486 cells, supporting our observations in cell line models and KRAS inhibitor-resistant cell lines (**Figures 4I-J, S4D**). We established an additional patient-derived cell line model of KRAS G12C lung cancer, DFCI516, using banked processed tumor material, and confirmed via Sanger sequencing that KRAS G12C was retained at high frequency as in DFCI486 (**Figure S4E**). Interestingly, live-cell imaging assays revealed that while DFCI486 and DFCI516 were intrinsically resistant to sotorasib, FHD-286 significantly sensitized cells to sotorasib (**Figure 4K-L**). Further, DFCI516 cells receiving drug combinations failed to regrow following drug discontinuation as compared to FHD-286 alone (**Figure 4L**). Further supporting our mechanistic understanding for this sensitization occurring via the EMT-axis, we achieved comparable responses with the combination of sotorasib and the AXL inhibitor, bemcentinib (**Figure S4F**), highlighting the essentiality of AXL signaling for the survival of DFCI516 and its required regulation by mSWI/SNF complexes. Together, our results implicate mSWI/SNF complexes as master regulators of treatment failure in KRAS-mutant NSCLC facilitated by dysregulation of AXL-mediated cellular processes which include survival and EMT.

### FHD-286 enhances *in vivo* anti-tumor efficacy of sotorasib treatment in patient-derived xenograft mouse models

Finally, we sought to define whether combined KRAS and mSWI/SNF inhibition resulted in meaningful impacts in tumor models *in vivo*. To this end, we evaluated KRAS G12C NSCLC adenocarcinoma xenograft-derived organoid (XDO) models^43^ encompassing a wide range of co-existing genomic alterations, and *in vivo* sensitivities to KRAS G12C inhibitor (G12Ci) monotherapy (**Figure S5A**). In both the XDO_PHLC344 (G12Ci-resistant) and XDO_PHLC207 (G12Ci-sensitive), FHD-286 alone had potent single-agent activity, with no added benefit from concurrent sotorasib treatment (**Figure S5B**). Interestingly, both the XDO_PHLC239 (G12Ci resistant) and XDO_PHLC194 (G12Ci-sensitive) organoid models showed additive activity from the combination of sotorasib and FHD-286 treatment (**Figure 5A**).

**Figure 5.**
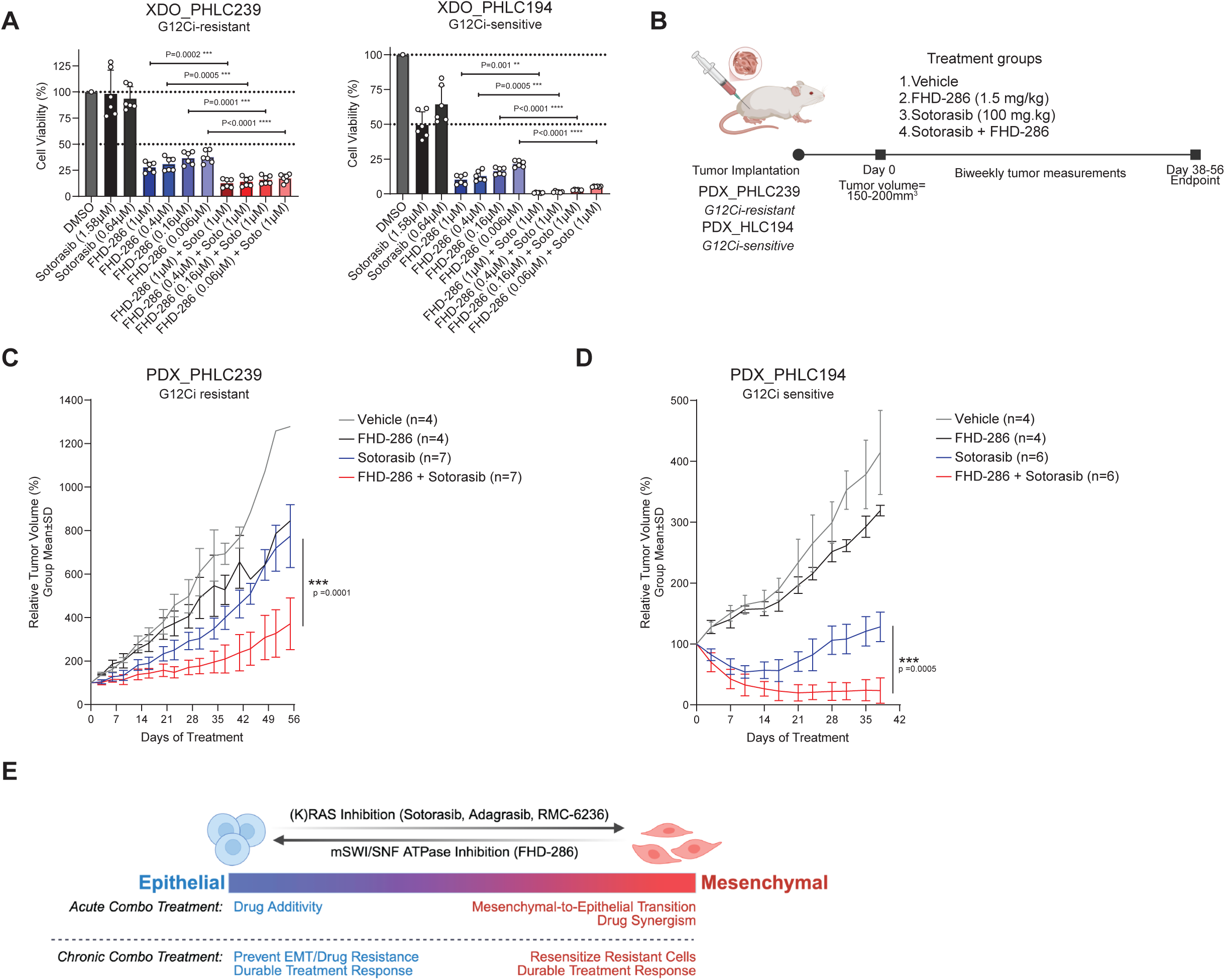
Combination FHD-286 and sotorasib treatment results in durable anti-tumor efficacy in G12Ci-resistant and -sensitive tumors in vivo. A. Patient-derived organoids, XDO_PHLC239 and XDO_PHLC194 embedded in Matrigel and treated according to conditions indicated. 3D cell titer Glo (CTG) measurements were performed on Day 14. B. Schematic of in vivo PDX experimental design. PDX tumor pieces were implanted into mouse flank and grown to 150-200mm3 before receiving indicated drug treatments. C-D. In vivo tumor growth kinetics of PDX models, PHLC239 (C) and PHLC194 (D), displayed as relative tumor volume and analyzed by linear mixed-effects modeling. p-values are indicated. E. Model describing response to KRAS and mSWI/SNF ATPase inhibition in NSCLC cells across a spectrum of EMT status.

Given these results, we next performed *in vivo* xenograft studies using patient-derived xenograft (PDX) models PHLC239 and PHLC194 (**Figure 5B**), representative of both G12Ci resistant and sensitive organoid models, respectively^44^ (**Figure S5A**). As PDX_PHLC239 was refractory to sotorasib treatment, only combination treatment significantly decreased tumor volume following 56 days of treatment with no significant adverse effects on animal weight (**Figures 5C, S5C-D**). Further, in our PDX_PHLC194 model, tumors were initially responsive to sotorasib monotherapy, with maximum tumor regressions achieved around day 14, after which acquired resistance to sotorasib was observed (**Figures 5D, S5C**). Importantly, concurrent FHD-286 treatment significantly decreased tumor growth kinetics compared to sotorasib alone (P<0.0001) and led to deeper and more sustained response following 39 days of treatment, again without affecting overall animal weight (**Figures 5D, S5C-D**). Together these results broaden our understanding of the therapeutic benefit of mSWI/SNF inhibition and reveal the efficacy of FHD-286 in re-resensitizing resistant KRAS G12C models to sotorasib treatment.

## Discussion

KRAS mutations are among the most common oncogenic drivers in human cancer, underscoring an urgent need for treatment approaches that effectively and durably inhibit KRAS signaling. Despite the recent emergence of mutant-selective and pan-RAS mutant inhibitors, to date, therapeutic responses in KRAS-mutant lung cancers remain inefficient and short-lived. Efficacy is often limited by adaptive mechanisms that reactivate downstream signaling or invoke alternative survival pathways ^1^. Here we identify mSWI/SNF (BAF) chromatin remodeling complexes as central regulators of KRAS inhibitor response and resistance. We reveal that the cellular plasticity of KRAS-mutant lung cancer cells along the epithelial-mesenchymal axis is shaped by mSWI/SNF-mediated chromatin accessibility and gene regulation. Importantly, our findings position the clinical-grade mSWI/SNF SMARCA4/2 ATPase inhibitor, FHD-286, as a potential strategy to circumvent adaptive resistance programs in KRAS-mutant lung cancers.

Our studies demonstrate that mSWI/SNF inhibition enhances KRAS inhibitor response through two convergent mechanisms: dampening EMT and AXL-driven bypass signaling, and attenuating proliferative programs through cell-cyle related pathways, which re-merge during the acquisition of KRAS inhibitor resistance. As such, drug synergy is most prominent in models with strong EMT signatures, whereas epithelial-leaning models tend to lack acute synergy but display enhanced durability of response stemming from blockade of EMT-mediated resistance (**Figure 5E**). In NSCLC, AXL has a well-documented role in EMT and has been linked to resistance to tyrosine kinase inhibitors (TKIs) ^30,45,46^. Our work extends previous findings from our group implicating mSWI/SNF complexes in TKI resistance in EGFR-mutant lung cancers via modulation of the NRF2-ROS axis ^11^, revealing here that in KRAS-mutant NSCLC, mSWI/SNF complexes can maintain a mesenchymal state and support a chromatin architecture permissive of adaptive regrowth after KRAS inhibition. This occurs uniquely through mSWI/SNF-mediated regulation of known pathways which dampen KRAS inhibitor efficiency and promote resistance, such as EMT, cytoskeletal remodeling, TNF/NF-κB, and TGF signaling, suggesting that complex inhibition works by undoing adaptive state changes that occur in cells to compensate for loss of upstream oncogenic signaling cues. Notably, recent studies have highlighted how lineage plasticity is a critical mechanism of non-genetic resistance to KRAS-targeted therapies in lung adenocarcinomas as well as in pancreatic and colorectal cancers^4,27,30,47^, reinforcing how chromatin remodeling can be a unifying, broadly relevant targetable axis of KRAS therapeutic failure.

Our results encourage classifying KRAS mutant tumors not solely on mutational subtype but also by expression of cell state-defining biomarkers. This is exemplified by our finding that synergy was independent of *STK11* mutational status, but highly dependent on EMT status. In STK11-mutant tumors, EMT together with NF-κB and interferon alpha signaling pathways may influence immune evasion, which is a key vulnerability in STK11-mutant KRAS tumors ^48–51^. With evidence that intrinsic cell of origin properties may identify tumors that undergo EMT from those which do not ^52^, determining whether the high EMT signature characterized in a subset of our model systems stems from differences in cell of origin may be valuable. Our findings thus underscore the importance of chromatin states in deciding tumor “transcriptional phenotypes” that predict therapeutic response. Future investigation into how chromatin state reprogramming could render immune-cold KRAS tumor environments amenable to immunotherapies may inform connectivity between mSWI/SNF function and immune modulation ^53–56^.

Combination mSWI/SNF and KRAS inhibition is broadly effective, spanning multiple KRAS mutant variant cell lines (G12C/D/A/S/V, G13D, Q61H), *ex vivo* patient-derived spheroids, XDOs, and *in vivo* PDX model systems. In KRAS-mutant models with either intrinsic or acquired resistance to sotorasib or adagrasib, mSWI/SNF inhibition was sufficient to reverse resistance and sensitize models to KRAS-targeted therapies. Because this was consistent across mutation-specific inhibitors (sotorasib, adagrasib, MRTX-1133) and the pan-RAS inhibitor RMC-6236, these results raise the intriguing possibility for small molecule ATPase inhibition to be used as a preconditioning strategy. Finally, combination therapy allowed for sustained suppression and prevention of bypass signaling even after drug withdrawal, suggesting that mSWI/SNF activity collapse may imprint a heritable chromatin state that constrains the future plasticity of cancer cells. This aligns with the known role of mSWI/SNF remodelers in promoting lineage-defining chromatin states^57–59^ via its ability to interact with lineage-specific transcription factors ^60^, establish chromatin memory retained through neoplastic transformation ^61^, or other mechanisms in which SWI/SNF remodelers maintain a malignant chromatin state^62^.

As mSWI/SNF complexes integrate cellular signals to dynamically regulate chromatin architectures supportive of specialized cellular functions, pharmacologic manipulation of these downstream nodes may offer advantages over other parallel inhibition strategies which usually become circumvented (eg. KRAS/EGFR, BRAF/MEK inhibitors)^63,64^. mSWI/SNF inhibition, thereby, may block adaptive resistance programs that are otherwise difficult to counteract with single pathway inhibitors, unlocking the unique cooperativity between kinase inhibitor-induced signaling shutdown and mSWI/SNF-dependent chromatin priming. Together, our findings support a therapeutic model in which targeting mSWI/SNF function not only enhances the depth and durability of KRAS inhibitor response, but also hinders the adaptive reprogramming that underlies targeted therapy resistance, providing strong mechanistic foundations for evaluating combination strategies in diverse tumor types.

## Acknowledgements

We thank members of the Kadoch, Jänne, and Sacher Laboratories for assistance and discussion throughout the duration of this project. We also thank Zach Herbert and Maura Sullivan of the DFCI Molecular Biology Core Facility (MBCF) for help with high-throughput sequencing. C.G. is supported by the Next Generation of Scientists (NGS) Program from the Cancer Research Society (Grant 1156629). W.W.F. is supported by a Lung Cancer Research Foundation grant (Award 1307903). FTHW is supported by the University of British Columbia Clinician Investigator Program. This work was supported in part by the National Cancer Institute (R35 CA220497; P.A.J.), and the American Cancer Society (CRP-17-111-01-CDD; P.A.J.). This work was supported in part by the Howard Hughes Medical Institute (HHMI), with the generous support of Roy and Carol Beerman and the Meredith and Billy Starr Investigatorship (C.K.). This work utilized an Illumina NovaSeq X Plus (DFCI) that was purchased with funding from a National Institutes of Health SIG grant 1S10OD036228-01.

## Author Contributions

CG, WWF, SL, and CK: Conceptualization, methodology, validation, formal analysis, investigation, data curation, writing, visualization, project administration. DCC and AWY: formal analysis, data visualization, methodology. FTHW, NAP, NR, KH: formal analysis, conceptualization - PDOs, manuscript editing. NAP, NR, PMC, QL: data curation. Joseph K: investigation, data curation, methodology, validation. MMH: data curation and methodology. LL, JAT: investigation, data curation and methodology. Jens K, FF, JL: data curation. CW, MAL, PCG: methodology. KN: investigation and methodology. MST, AGS, PAJ, CK: supervision, resources and funding acquisition.

## Declaration of Interests

A.G.S: Institutional Research & Clinical Trial PI: AstraZeneca, Amgen, Genentech, Merck, Lilly, Pfizer, BMS, Spectrum, GSK, Iovance, CRISPR Therapeutics, BridgeBio, HotSpot Therapeutics, AdaptImmune. Advisory committee (no personal fees): Genentech, Amgen, Merck. Travel expenses for clinical trial investigator meetings: Amgen, Merck, Genentech-Roche.

P.A.J. has served as a consultant for AstraZeneca, Boehringer Ingelheim, Pfizer, Roche/Genentech, Chugai Pharmaceuticals, Eli Lilly pharmaceuticals, SFJ Pharmarceuticals, Voronoi, Daiichi Sankyo, Biocartis, Novartis, Sanofi,Takeda Oncology, Mirati Therapeutics, Transcenta, Silicon Therapeutics, Syndax, Nuvalent, Bayer, Eisai, Allorion Therapeutics, Accutar Biotech, Abbvie, Monte Rosa Therapeutics, Scorpion Therapeutics, Merus, Frontier Medicines, Hongyun Biotechnology, Duality Biologics, Blueprint Medicines, Dizal Pharma, GlaxoSmithKline, Tolremo, Myris Therapeutics, Bristol Myers Squibb. P.A.J. has received research funding from AstraZenenca, Daiichi Sankyo, PUMA, Eli Lilly pharmaceuticals, Boehringer Ingelheim, Revolution Medicines, and Takeda Oncology. P.A.J. receives-marketing royalties from Dana Farber Cancer Institute owned intellectual property on EGFR mutations licensed to Lab Corp. C.K. is the Scientific Founder, Scientific Advisor to the Board of Directors, Scientific Advisory Board member, shareholder, and consultant for Foghorn Therapeutics, Inc. (Cambridge, MA), and serves on the Scientific Advisory Board of Nereid Therapeutics. C.K. is also a member of the *Molecular Cell* and *Cell Chemical Biology* Editorial Boards. F.F. has received personal fees from Roche for speaker activity. The other authors declare no competing interests.

## Materials and Methods

### Cell lines

H2030, H2122, H358, H1373, LU-65, HOP62, HCC44, H23, A549, H2009, H1944, H441, HCC461, and DFCI316 cells were maintained in RPMI-1640 (Gibco) supplemented with 10% fetal bovine serum (GeminiBio) and 1% penicillin-streptomycin (Gibco). The identities of commercial cell lines were verified by STR testing (Dana-Farber Cancer Institute). DFCI516 and DFCI486 patient-derived cell lines were newly established using our previously described methodology (Kohler et al MCT 2021). All patients were provided written informed consent. All studies were conducted under a Dana-Farber Cancer Institute IRB approved protocol and in accordance with the Declaration of Helsinki. All cell lines were routinely monitored for mycoplasma contamination and confirmed negative by PCR analysis (ATCC).

### Generation of resistant cell lines

H358SR and H358AR cells were generated by culturing H358 parental cells in 100 nM sotorasib and 1000 nM adagrasib, respectively, for >4 months. Cells were maintained in RPMI-1640 (Gibco) supplemented with 10% fetal bovine serum (GeminiBio), 1% penicillin-streptomycin (Gibco), and their respective drugs.

### Western blotting

Whole cell and nuclear lysates were generated either using RIPA lysis buffer (50 mmol/L Tris, pH 8.0, 150 mmol/L NaCl, 5 mmol/L MgCl2, 1% Triton X-100, 0.5% sodium deoxycholate, 0.1% SDS) or using no salt EB0 buffer (50mM Tris, 0.1% NP-40, 1mM EDTA, 1mM MgCl_2_) followed by high salt EB300 buffer (50mM Tris, 1% NP-40, 1mM EDTA, 1mM MgCl2, 300mM NaCl), each supplemented with protease and phosphatase inhibitor cocktails (Thermo Fisher, Cat#78440), respectively. Equal amounts of total protein were separated by SDS-PAGE and blots were probed as indicated (all antibodies are found in KRT). Signals were detected using either SuperSignal West Pico PLUS (Thermo Fisher), Femto chemiluminescent substrates (Thermo Fisher), and imaged on an Amersham Imager 600.

### Cell viability and Synergy assays

Dose-response curves and cell viability assays were performed by seeding 500 cells per well into 384-well plates. Cells were drugged the following morning in triplicate. Cell viability was assessed after 96 hours with CellTiter-Glo Luminescent Cell Viability Assay (Promega) according to manufacturer’s instructions. Plates were read using a POLARstar Omega microplate reader (BMG Labtech). IC50 values were calculated using the following formula: normalized viability = 100/(1+ (inhibitor)/IC50)^HillSlope in GraphPad Prism v10.2.1.

Drug synergy was assessed with Combenefit software as previously described ^24^. Cells were drugged in 6-by-6 drug concentration matrices in triplicate and viability was measured by CellTiter-Glo after 96 hours. Loewe synergy scores were calculated for each drug combination and were mapped relative to cell proliferation using Combenefit v2.021.

### Live-cell imaging growth assays

96-well plates were housed in a BioSpa 8 automated incubator (Agilent) and were scanned at regular intervals using a Cytation5 cell imaging multimode reader (Agilent) or were housed and scanned with a CELLCYTE X (Cytena). Media was replenished approximately every 4 days. Image analysis was conducted using Gen5 v3.15 and CELLCYTE Studio v2.7.4.

### Cloning and Lentiviral infection

EZ-Tet-pLKO-Hygro was a gift from Cindy Miranti (Addgene plasmid #85972; http://n2t.net/addgene:85972 ; RRID: Addgene_85972). We replaced the hygromycin resistance cassette was with a Blasticidin resistance cassette using In-Fusion HD cloning according to manufacturer’s instructions, resulting the Tet-pLKO_Blast backbone used for all dox-inducible knockdown studies in the manuscript. shRNA sequences targeting SMARCA2 were derived from the Broad Institute shRNA database (See **Table S3** for target sequences) and a control shRNA sequence was found from Addgene plasmid #1864. Oligonucleotides containing shRNA sequences were ordered from IDT and were annealed in Nuclease-Free Duplex Buffer (IDT) by heating to 100°C and slowly cooling to room temperature. Tet-pLKO_Blast was restriction digested with NheI-HF and EcoRI-HF for 1 hour at 37°C. Linearized vector was gel purified and duplexed shRNA oligonucleotides were ligated using standard T4 ligation. Ligation reaction was transformed into Stbl3 and colonies were screened with colony PCR to identify successful recombinants. Sequences of plasmids were confirmed with whole plasmid sequencing (Plasmidsaurus) before use.

lentiCRISPR v2 was a gift from Feng Zhang (Addgene plasmid # 52961 ; http://n2t.net/addgene:52961; RRID: Addgene_52961). CRISPR knockout gRNA sequences were derived from the Brunello genome-wide knockout library^65^ (See **Table S3** for target sequences). Oligonucleotides containing target sequences were ordered from IDT and were annealed in water by heating to 100°C and slowly cooling to room temperature. LentiCRISPRv2 was restriction digested with BsmBI at 37°C for 1 hour. Linearized vector was gel purified and duplexed shRNA oligonucleotides were ligated using standard T4 ligation. Ligation reaction was transformed into Stbl3 and colonies were screened with colony PCR to identify successful recombinants. Sequences of plasmids were confirmed with whole plasmid sequencing (Plasmidsaurus) before use.

pCW57 was a gift from David Root (Addgene plasmid # 41393 ; http://n2t.net/addgene:41393; RRID: Addgene_41393). cDNA sequences for STK11 were cloned from HCC4006 cells and AXL was cloned from HOP62. pCW57 was digested with BmtI-HF and AgeI-HF at 37°C for 1 hour. cDNA sequences were shuttled into pCW57 by InFusion HD cloning (Takara). Sequences of plasmids were confirmed with whole plasmid sequencing (Plasmidsaurus) before use.

Lentiviral particles were generated by transfecting lentiviral expression plasmid, psPAX2, and pMD2.G into HEK293T cells with FuGene HD (Promega). Lentivirus was harvested 48 hours after transfection and was passed through a 0.45 um syringe filter.

### Fluorescent Ubiquitination-based Cell Cycle Indicator (FUCCI) reporter

tFucci(CA)2/pCSII-EF was a generous gift from Hiroyuki Miyoshi and Atsushi Miyawaki ^29^. An IRES-hygromycin resistance cassette was ordered as a gBlock form IDT and was introduced by In-Fusion HD Cloning to allow for antibiotic selection of stable cells following lentivirus-mediated integration of the reporter construct. Cells were stably transduced with tFucci(CA)2/pCSII-EF in the presence of 10 µg/mL polybrene. Stable integrants were selected with 14 days of 500-800 µg/mL hygromycin treatment.

Cells were harvested following the indicated treatment conditions and were fixed in 4% PFA for 15 minutes at room temperature. Cells were washed in PBS and were stored in 4°C until all samples in the time course were collected. Percentage of cells in G1, S, or G2/M phase were measured as %mCherry+/mVenus-, %mCherry-/mVenus+, or %mCherry+/mVenus+, respectively, by flow cytometry on an LSRFortessa (BD). Data was analyzed with FlowJo v7.

### Whole Exome Sequencing

Genomic DNA was extracted using the DNeasy Blood & Tissue kit (Qiagen, Cat#69504) and was submitted for WES at the Broad Institute of MIT and Harvard using the ‘‘Express Somatic Human WES v6’’ workflow. Captured fragments were sequenced on a NovaSeq 6000 with an S4 flow cell according to standard Illumina protocols using 151 bp paired-end sequencing reads and achieved a median target coverage of ∼200X. Sequencing reads were aligned to human genome build 37 (GRCh37/hg19), aggregated into a BAM file, and further processed to identify somatic variants with GATK v4.0.4.0, MuTect2, and Oncotator v1.9.8.0. A panel of normal tissue samples were used to filter out germline variants, identify sequencing artifacts, and to serve as a reference for CNV analysis.

### Transwell Migration Assay

H2030 cells were treated with DMSO, 100 nM sotorasib, and/or 100 nM FHD-286 in RPMI-1640 media supplemented with 10% FBS for 6 days. Cells were serum starved overnight in the presence of drug before 100,000 cells in FBS-free media were seeded into 6.5 mm Transwell inserts with 8.0 µm pores (Corning 3464) the following morning. Transwell inserts were placed into wells of 24-well plates containing RPMI-1640 supplemented with 10% FBS. Both media inside and outside the insert contained drugs at the indicated concentrations. 72 hours after seeding, Transwell inserts were scrubbed with cotton tipped swabs to remove cells that failed to migrate. Cells on the underside of the membrane were fixed in methanol and stained in 0.5% Crystal Violet staining solution for 30 minutes. Inserts were washed in water twice before drying overnight. The following morning, the inserts were imaged on a Cytation5 Cell Imaging Multimode Reader (Agilent) and the number of cells per field were quantified using Gen5 v3.15 (Agilent).

Representative images are shown for a single experiment from N=2 independent experiments. The bar graph depicts the number of cells that migrated through the Transwell membrane in three distinct fields of view from two biological replicates per treatment condition. 0% FBS controls were included in which untreated cells were seeded into FBS-free media in Transwell inserts and placed into wells with RPMI-1640 without FBS supplementation to account for migration occurring in the absence of a nutrient gradient. Statistical significance between groups was assessed by one-way ANOVA with Sidak’s multiple comparisons test. *** p < 0.0001.

### CUT&RUN

CUT&RUN was performed as previously described ^11^ using Epicypher protocol for their CUTANA ChIC/CUT&RUN kit. 500,000 cells were harvested per antibody in each condition per cell line evaluated and washed with PBS before proceeding with protocol. To note, a nuclear extraction step using NE buffer (20 mM HEPES–KOH pH 7.9, 10 mM KCl, 0.1% Triton X-100, 20% Glycerol supplemented with fresh 0.5 mM Spermidine and 1X protease inhibitor) was performed on all cell lines except H2030, H2122 (as these cell lines were not amenable to this step for successful immunoprecipitation). Sample purification was done using SPRIselect reagent beads at 1x ratio at end of day 1. CUT&RUN sample library preparation was done using Epicypher Library Prep kit, using manufacturer’s protocol with a slight modification. SPRIselect reagent bead purification following PCR amplification was performed at a 0.8X ration of beads to DNA to efficiently remove primer dimers. Libraries were analyzed on Tapestation (Agilent) and quantified on Qubit before paired-end sequencing on NovaSeqX Plus (DFCI).

### ATAC-seq

The Omni-ATAC protocol ^66^ was used with slight modifications as previously described ^11^. 100,000 cells per condition were used and washed with PBS before cell lysis (10mM Tris–HCl pH 7.4, 10mM NaCl, 3mM MgCl2, NP-40 (final 0.1% v/v), Tween-20 (final 0.1% v/v) and digitonin (final 0.01% v/v)) for 3 minutes on ice. The Qiagen MinElute Reaction Cleanup Kit was used for DNA purification and library preparation was done using PCR amplification. ATAC-seq libraries were sequenced on NovaSeq 6000 (DFCI) using 37-bp paired-end sequencing. ATAC-seq experiments were performed in biological duplicates for all cell lines and experimental conditions.

### RNA-seq

Half a million to one million cells were harvested, washed 1X with cold PBS, resuspended in 350ul RLT buffer (Qiagen) and stored at -80C until further processing. Sample was passed through the QIAshredder (Qiagen) system and RNA was purified using the Qiagen RNeasy kit (Qiagen) and further processed using the Illumina NEBNext Ultra II Directional RNA Library Prep Kit (New England Biolabs). Library quality control was assessed by TapeStation (Agilent) and were quantified by Qubit Fluorometer. RNA libraries were sequenced on the Illumina NextSeq 500 (DFCI) with 75-bp single-end sequencing. RNA-seq experiments were performed in biological duplicates or triplicates for all cell lines and experimental conditions as reported in figure legends.

### *Ex vivo* tumor spheroid drug screens

DFCI486 and DFCI491 patient-derived xenografts were expanded in immune-deficient NOD SCID gamma (NSG) mice. Tumors were harvested and processed according to our previously published protocols ^41,42^. Briefly, tumors were minced using scalpels and were digested in pre-warmed RPMI-1640 supplemented with an enzyme cocktail consisting of collagenase, dispase, DNase I, and ROCK inhibitor at 37°C for 20 minutes. Digested material was serially filtered through 100 µm and 40 µm cell strainers. Cellular contents between 40-100 µm were seeded in 384-well black-wall clear bottom plates and immediately drugged using an HP D300 Digital Dispenser. CellTiter-Glo 3D viability assay was run at 96 hours post-treatment.

### *In vitro* organoid drug screens

This study uses 4 human lung adenocarcinoma xenograft-derived organoids (XDO), XDO_PHLC344, XDO_PHLC239, XDO_PHLC194 and XDO_PHLC207, derived from the genomically annotated PDX cohort as previously reported by Mirhadi et al ^43^ and maintained by the Princess Maragaret Living Biobank core facility using previously described protocols ^67^. XDO-models were authenticated by short tandem repeat analysis as previously described^68^. For the XDO-models, *in vitro* drug efficacy studies were done in technical triplicates. Organoids were dissociated into single-cell suspensions and plated into Matrigel-coated 96-well plates (5,000 cells per well, suspended in a 30μL slurry of 50% Matrigel plus 50% of M26 growth media^68^). The plates were gently spun down to allow the slurry to solidify into domes around cell pellets over 5-10min, then overlaid with 100μL of M26 media per well, and kept in 37°C 5% CO_2_. Twenty-four hours later, drugs were dispensed into each well at specified concentrations (spanning the range of 0.001 to 100 μmol/L) using a Tecan D300e Digital Dispenser. Sotorasib was purchased from UHN-Shanghai and reconstituted in DMSO, FHD-286 was provided by the Kadoch laboratory and reconstituted in DMSO. Media was changed every 3-4 days with fresh drugs, maintaining the same initial concentrations. Fourteen days later, cell viability was determined using the CellTiter-Glo® 3D viability assay. Drug-response curves were plotted using GraphPad Prism 10.0 software.

### *In vivo* xenograft studies

Patient-derived xenografts (PDXs) were generated from primary tumor tissue collected from patients undergoing surgical resection of early-stage non-small cell lung carcinoma (NSCLC) tissue, according to previously published protocols^69^(10.1158/1078-0432.CCR-10-2224), with approval from the University Health Network Ethics Board (REB #09-0510-T) with informed written consent from all participating patients, and the Animal Care Committee (AUP #5555). These studies were all conducted in accordance with the Canadian Tri-Council Policy Statement: Ethical Conduct for Research Involving Humans – TCPS 2 (2022).

For each of the selected KRAS^G12C^ NSCLC adenocarcinoma PDX models, cryopreserved PDX tissue (below passage 10) was thawed and implanted subcutaneously into the flank of a NOD-SCID male mouse. Tumors were harvested at endpoint volume and cut into 3-mm diameter pieces for expansion into sufficient sample size (n∼20 mice) to populate the required experimental arms. Once average tumor volumes reached 150-200mm^3^, mice were stratified equally by tumor volumes and weights into treatment groups for therapy initiation by daily oral gavage. Treatments were either vehicle 1 (5% DMSO + 40% PEG300 + 5%Tween 80 + 50% MilliQ-H2O), FHD-286 (SMARCA4/2 inhibitor; 1.5mg/kg; Kadoch laboratory) in vehicle 1, and sotorasib (100 mg/kg; UHN-Shanghai, Shanghai, China) in vehicle 2 (0.5% methylcellulose + 1% Tween-80), or the combination given sotorasib first, followed by FHD-286 with at least 1 hour in between. Tumor sizes (width x length) were measured by caliper twice weekly until humane endpoints were reached.

Tumor growth kinetics were then analyzed by linear mixed-effects regression. Absolute tumor volumes (TVs) measured serially over time were normalized to day zero measurements to derive relative TVs. Independent variables included fixed effects for time (measured in weeks), treatment group (either sotorasib or sotorasib+FHD-286), and the interaction between time and treatment. A random intercept for each animal ID was also included. For each model, all measurements were included from day zero until the last date when both groups had at least one mouse still being followed. A statistically significant interaction between time and treatment indicated different growth rates between treatment groups.

### Data and code availability

All sequencing raw and processed data have been deposited in the Gene Expression Omnibus (GEO) database and are publicly available upon publication under the series GSE313345.This paper does not report original code.

### Assessing BAF and kinase mutational and gene expression patterns in primary cancer samples

Using data from The Cancer Genome Atlas (TCGA) program (TCGA Research Network; https://www.cancer.gov/tcga), we assessed patterns of variation in BAF subunit and kinase genes. MAF files for TCGA lung adenocarcinoma (LUAD) were collected from CBioPortal ^70–72^ and maftools v. 2.17.10 ^73^ was used to filter away the top 10% most mutated samples, which were considered “hyper mutated”, and to generate an oncoplot summarizing the prevalence and identity of variation across genes. Variation in *KRAS* among the LUAD cohort was summarized using a pie chart constructed using ggplot2 v. 3.5.1.

TCGA LUAD variation data were used to gather a set of samples with the following profiles: (1) all samples with *KRAS* variation (N = 144), (2) samples with *KRAS* variation and specific patterns of variation in the BAF subunit genes (N = 33), and (3) samples with *KRAS* variation and no specific patterns of variation in the BAF subunit genes (i.e., “WT” status; N = 111). The specific patterns of variation in the BAF subunit genes used to stratify samples were nonsense or frameshift variants or missense variants that fell within the ARID/CBR domains of *ARID1A* or the ATPase domain of *SMARCA4*.

For each of the three sets of samples above that differ in the status of *KRAS* or BAF (*ARID1A* or *SMARCA4*) variation, we gathered raw expression count data for these samples and a similar number of “normal” lung samples from GTEx ^74^ using the recount3 resource ^75^, which ensures that the data were uniformly processed. We used DESeq2 v. 1.42.1 ^76^ to assess differential expression between each set of “normal” GTEx samples and corresponding *KRAS* or *BAF*-mutant samples. We excluded genes from the analysis if the raw count fell below 10 for at least half of the samples in the dataset. Ingenuity Pathway Analysis (IPA) was used to conduct an upstream regulator enrichment analysis based on log2(fold change) thresholds of -2 down and 2 up and an FDR p-value <= 0.05. We visualized the ranks of top-ranked upstream regulators across these three comparisons using the ggbump v. 0.1.0 package in R v. 4.3.1 ^77^. We also visualized the relative ranks of various epigenomic complex genes across these three comparisons using a table with the size and color scaled by rank.

### Assessing BAF and kinase mutational and gene expression patterns in cancer cell lines

DepMap Public 24Q4 data were downloaded from the DepMap portal (https://depmap.org/portal/)^16^ and were summarized in much the same fashion as the TCGA LUAD data outlined above. Briefly, maftools was used to produce an oncoplot based on the data contained in the omicsSomaticMutationsProfile MAF dataset. Of the DepMap LUAD cell lines with corresponding RNA-seq expression data: (1) 29 cell lines contained variation in *KRAS* overall, (2) 15 cell lines contained *KRAS* variation but no BAF subunit variation, and (3) 14 cell lines contained *KRAS* variation and BAF subunit variation. Again, we defined BAF subunit variation as nonsense or frameshift variants or missense variants that fell within the ARID/CBR domains of *ARID1A* or the ATPase domain of *SMARCA4*. Each of these three sets of cell lines was compared to a set of 6 DepMap “normal” lung control cell lines – PR-73nc9C, PR-bNFpIV, PR-BSHttf, PR-CPi0aO, PR-pVxTN4, and PR-VG2HGb – to assess differential expression using DESeq2. The results were anlyzed using IPA to identify enriched upstream regulators and their ranks across datasets were summarized as described above for the TCGA LUAD data.

### Assessing CCLE susceptibility to FHD-286

DepMap PRISM data were downloaded from the Skyros DepMap portal (https://theprismlab.org/portal/projects) (see **Table S1**). We specifically isolated drug sensitivity screen data for FHD-286 for LUAD cell lines with *KRAS* variation (N = 24; N = 31 total *KRAS*-variant cell lines) and visualized the effect of drug treatment using the area under the curve (AUC) values.

### Defining EMT score

EMT score was calculated using the K-S method previously described for RNA-seq transcriptomic data ^25,26^. Probe names were converted to HGNC symbols using DAVID^78,79^.

### Data processing for RNA-seq, ATAC-seq, and CUT&RUN

The O2 High Performance Compute Cluster, which is supported by the Research Computing Group at Harvard Medial School, was used for most data processing. See more information at https://it.hms.harvard.edu/about/departments/research-computing.

For a subset of datasets, sequencing was performed in batches and the raw FASTQ read data for these batches were combined for each unique sample.STAR v. 2.7.9a ^80^ with the options “--quantMode GeneCounts” was used to map raw RNA-seq reds to the human genome assembly hg19 with the refFlat annotation. Raw gene counts from the last column of the GeneReadsOut.tab file for each sample were gathered into a count matrix. RPKM normalization was applied to the raw counts using the hg19 refFlat annotation with the median isoform length for each gene.

Trimmomatic v. 0.36 ^81^ with the settings “ILLUMINACLIP:TruSeq3-PE.fa:2:30:10 CROP:30” was used to quality trim the raw ATAC-seq and CUT&RUN reads. These quality trimmed reads were mapped to the human genome (hg19 assembly) using Bowtie 2 v. 2.5.1 ^82,83^ with default settings except “-X 2000”. Picard MarkDuplicates v. 2.27.5 was used to filter away PCR duplicates. Narrow peaks were identified for CUT&RUN data using MACS2 v. 2.1.1.20160309 ^84^ with the settings “--nomodel -q 0.001”. Broad peaks were called for ATAC-seq data using MACS2 with the settings “-f BAMPE --broad --broad-cutoff 0.05 --nomodel -q 0.001”. Bedtools v. 2.31.1 ^85^ was used to create reference peak sets for quantitatively assessing accessibility (ATAC-seq) or occupancy (CUT&RUN) by merging peaks called across conditions for each dataset and for each antibody mark assayed using CUT&RUN. These reference peak sets and the bedtools intersect tool (with default settings) were used to gather raw counts of accessibility (ATAC-seq) or occupancy (CUT&RUN). Total sample library read coverage and peak widths from the appropriate reference peak set were used to normalize occupancy counts via RPKM normalization. The deepTools v. 3.5.0 ^86^ bamCoverage tool with settings “--binSize 40 --normalizeUsing CPM --exactScaling” was used to generate tracks of read coverage (i.e., bigwig files) for data visualization, which was conducted in the integrated genomics viewer (IGV)^87^.

### Analysis of RNA-seq, ATAC-seq, and CUT&RUN

R v. 4.3.1 was generally used for statistical analyses and data visualization. The R package eulerr v. 7.0.2 ^88^ was used to create Euler diagrams summarizing overlaps of raw peaks. PCA analyses performed on RPKM counts were used to confirm clustering of experimental replicates prior to downstream analyses.

DESeq2 v. 1.42.1 was used to assess differential expression between cell line replicates in comparisons between synergy and non-synergy cell lines or between experimental replicates in comparisons of drug treatments separately in each cell line (Sotorasib, FHD-286, and combination Sotorasib and FHD-286). Similarly, DESeq2 was used to assess differential accessibility between cell line replicates in comparisons between synergy and non-synergy cell lines or between experimental replicates in comparisons of drug treatments separately in each cell line (Sotorasib, FHD-286, and combination Sotorasib and FHD-286). Genes or ATAC-seq loci with a | fold change | >= 2 and FDR p-value <= 0.05 were inferred to be differentially expressed or accessible. Changes in occupancy of BRG1-SMARCA4, ARID1A, and H3K27ac based on replicate cell lines in each mark were assessed using edgeR v. 4.0.16 ^89^ with glmQLFit (apeglm v. 1.24.0)^90^ and those loci with a | fold change | >= 2 were called differentially occupied. The R package ggplot2 was used to generate MA and volcano plots for visualizing the results of these differential analyses. Bedtools and the hg19 refFlat gene annotation was used to identify the nearest gene TSS to each ATAC-seq or CUT&RUN locus and to calculate the distance to these features, which were summarized as barplots using ggplot2.

Functional enrichment of differentially expressed genes was evaluated using Metascape ^91^ or gene set enrichment analysis (GSEA)^92^. For GSEA, genes were ranked using log2(fold change) between experimental comparisons and clusterProfiler v. 4.10.1 ^93^ was used to run GSEA with a minimum gene set size of 25, a maximum gene set size of 500, and FDR correction for multiple statistical tests. MSigDB ^94^ gene sets for Hallmark were used for GSEA and the results were visualized as bubble plots of normalized enrichment scores (NES) and FDR p-values using ggplot2. Motif enrichment of cis-regulatory loci was assessed using HOMER v. 4.11 ^95^ with the findMotifsGenome.pl tool and a locus size of 400.

Heatmaps summarizing spatial patterns of accessibility or occupancy around peak loci (based on CPM-normalized signal) were generated using the deepTools tools computeMatrix (reference-point) and plotHeatmap with the options “--binSize 50 --missingDataAsZero”. ComplexHeatmap v. 2.18.0 ^96^ or ggplot2 were used to generate heatmaps of per-locus expression, accessibility, or occupancy. Row normalization of expression, accessibility, or occupancy was calculated using z-scores, which were colored using a blue to red color palette. Fold change measures between conditions were also visualized using a blue to red color palette.

---

**Figure S1.**
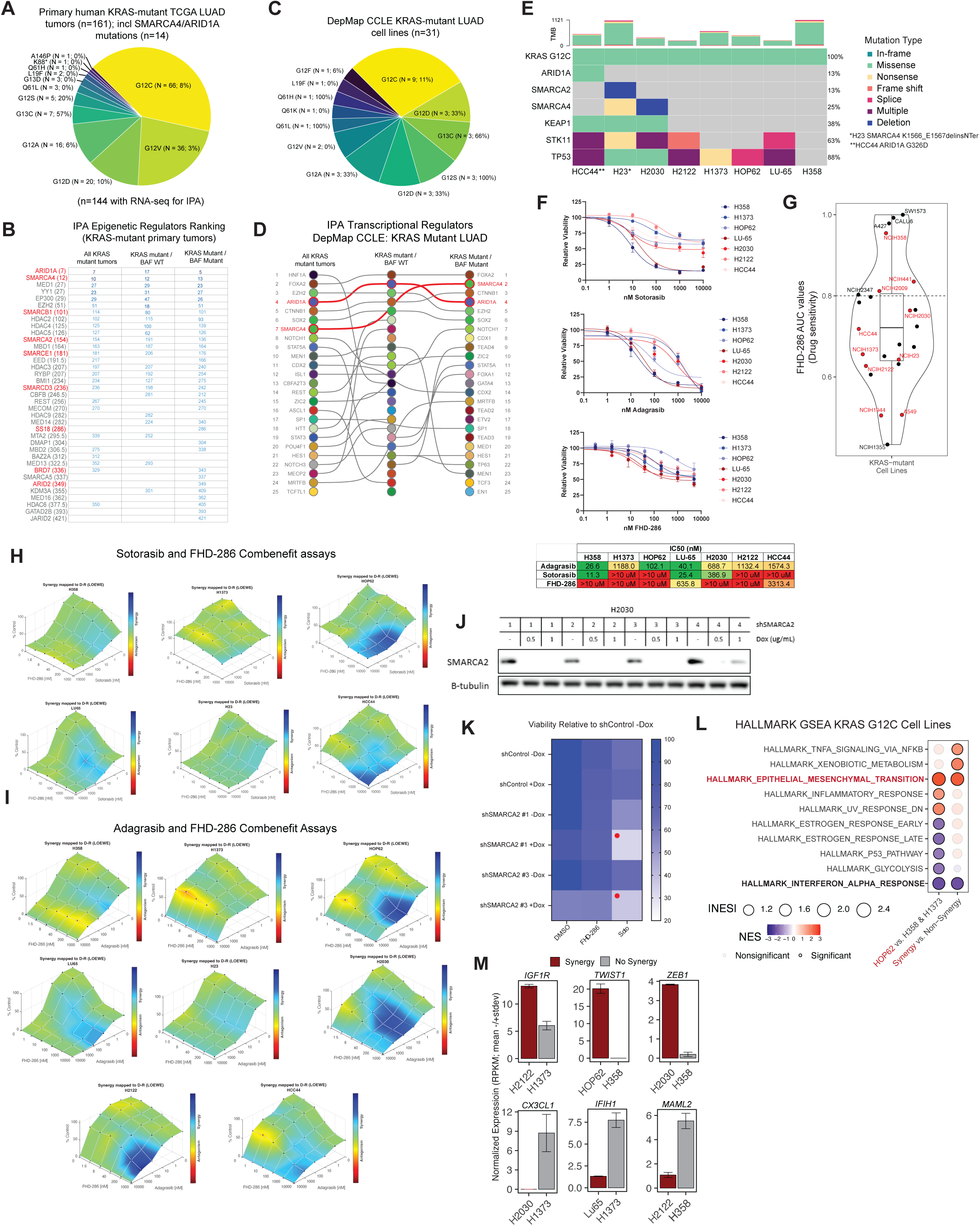
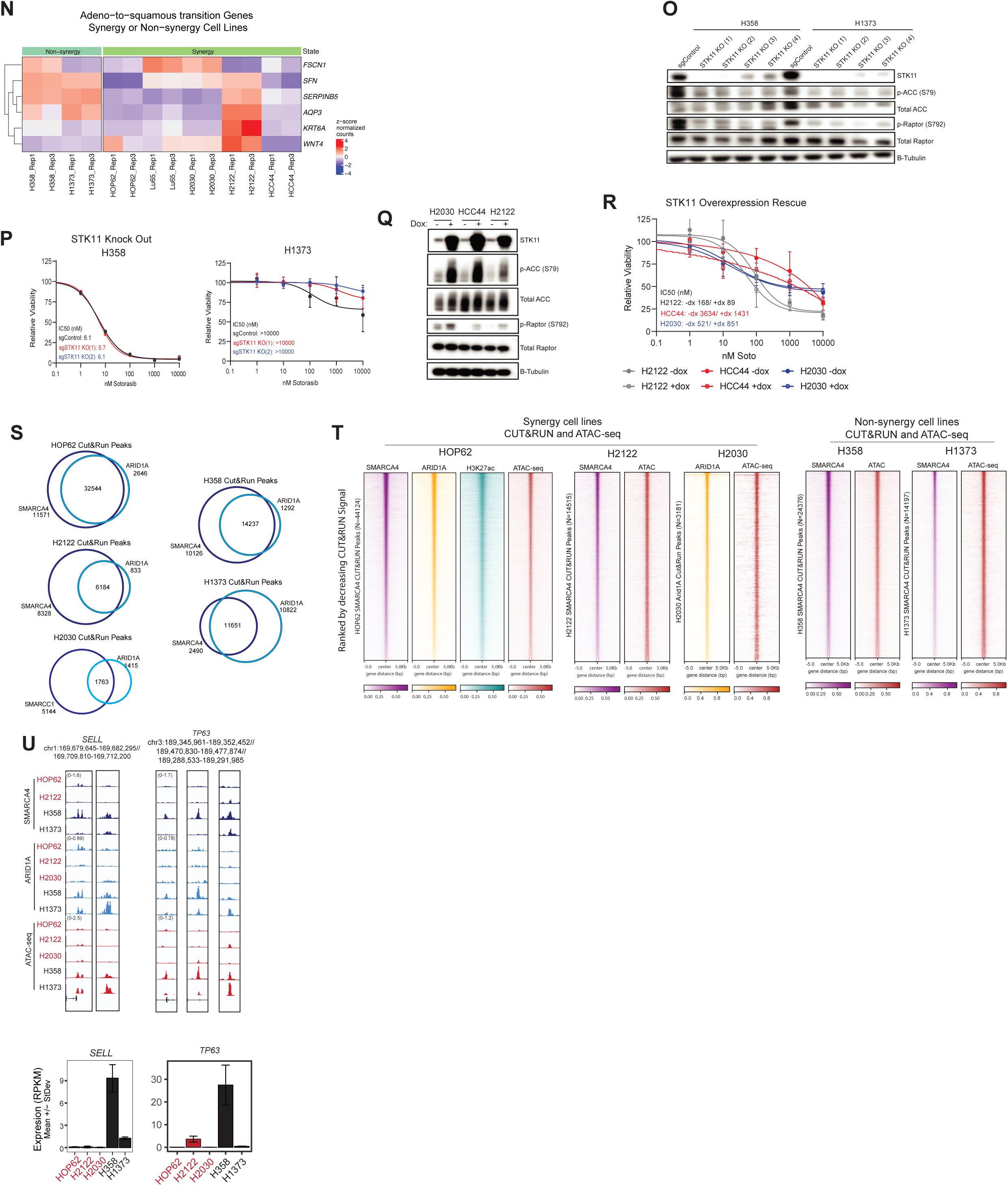
KRAS G12C-mutant cell lines are distinguished by specific mSWI/SNF chromatin features. A. Pie chart of KRAS mutations in TCGA LUAD primary tumors (total n=161, n=142 used in Fig1A IPA analysis). Samples include those with ARID1A and SMARCA4 mutations. B. Summary of ranks for epigenetic upstream transcriptional regulators (IPA) in KRAS-mutant LUAD primary tumors with and without BAF mutations; mSWI/SNF factors are highlighted. C. Pie chart of KRAS mutation type in CCLE cell lines (n=31). D. Results of upstream transcriptional regulator analysis in CCLE cell lines, analyzed by BAF mutation status. E. Mutation status of ARID1A, SMARCA4, STK11, KEAP1, and TP53 in KRAS-mutant cell lines used in this study. F. IC50 curves of sotorasib, adagrasib, and FHD-286 treatments across cell lines indicated. Table of IC50 values shown below. G. PRISM (Broad Institute) FHD-286 drug sensitivity AUC values for KRAS mutant cell lines; cell lines used in this study are shown in red; AUC>0.8 denotes FHD-286 sensitive lines (see also Table S1). H-I. Combenefit synergy studies performed at t=96 hours using (I) sotorasib and (J) adagrasib in combination with FHD-286. J. Immunoblot of SMARCA2 levels following dox induction of SMARCA2 shRNAs in H2030 cells. B-tubulin is loading control. K. Viabililty plot of SMARCA2 shRNA H2030 cells following DMSO, FHD-286 or sotorasib treatment. Viability measured relative to shControl with no dox treatment. Substantial decreases in cell viability upon SMARCA2 knock down and sotorasib treatment are indicated by red dots. L. Hallmark GSEA performed on HOP62 versus H358 and H1373 cell lines and all synergy versus non-synergy cell lines. M. Bar graphs indicating normalized expression values (RPKM) for top synergy- or non-synergy-specific genes. N. Gene signature for adenocarcinoma-to-squamous cell carcinoma in KRAS-mutant cell lines (REF) using biological replicates of RNA-seq. O. Immunoblot of STK11, phosphorylated and total ACC, phosphorylated and total RAPTOR levels upon knock out (KO) of STK11 in H358 and H1373 cells. B-tubulin is loading control. P. IC50 curves for sotorasib of H358 and H1373 cells upon STK11 KO. Q. Immunoblot of STK11, phosphorylated and total ACC, phosphorylated and total RAPTOR levels upon dox induction of STK11 over-expression in H2030, HCC44 and H2122 cells. B-tubulin is loading control. R. IC50 curves for sotorasib of respective cell lines upon over-expression of STK11. S. Venn diagrams showing overlap between mSWI/SNF CUT&RUN peaks for SMARCA4 or SMARCC1 and ARID1A across all cell lines. T. Heatmaps of SMARCA4 or ARID1A (CUT&RUN) occupancy and ATAC-seq in synergy and non-synergy cell lines. H3K27ac CUT&RUN correlates with BAF occupancy and ATAC signal in HOP62 cells. U. Top, Representative tracks at the SELL, and TP63 loci showing mSWI/SNF complex occupancy (ARID1A and SMARCA4 CUT&RUN) and ATAC-seq; bottom, RNA-seq normalized expression (RPKM) bar plots of SELL and TP63.

**Figure S2.**
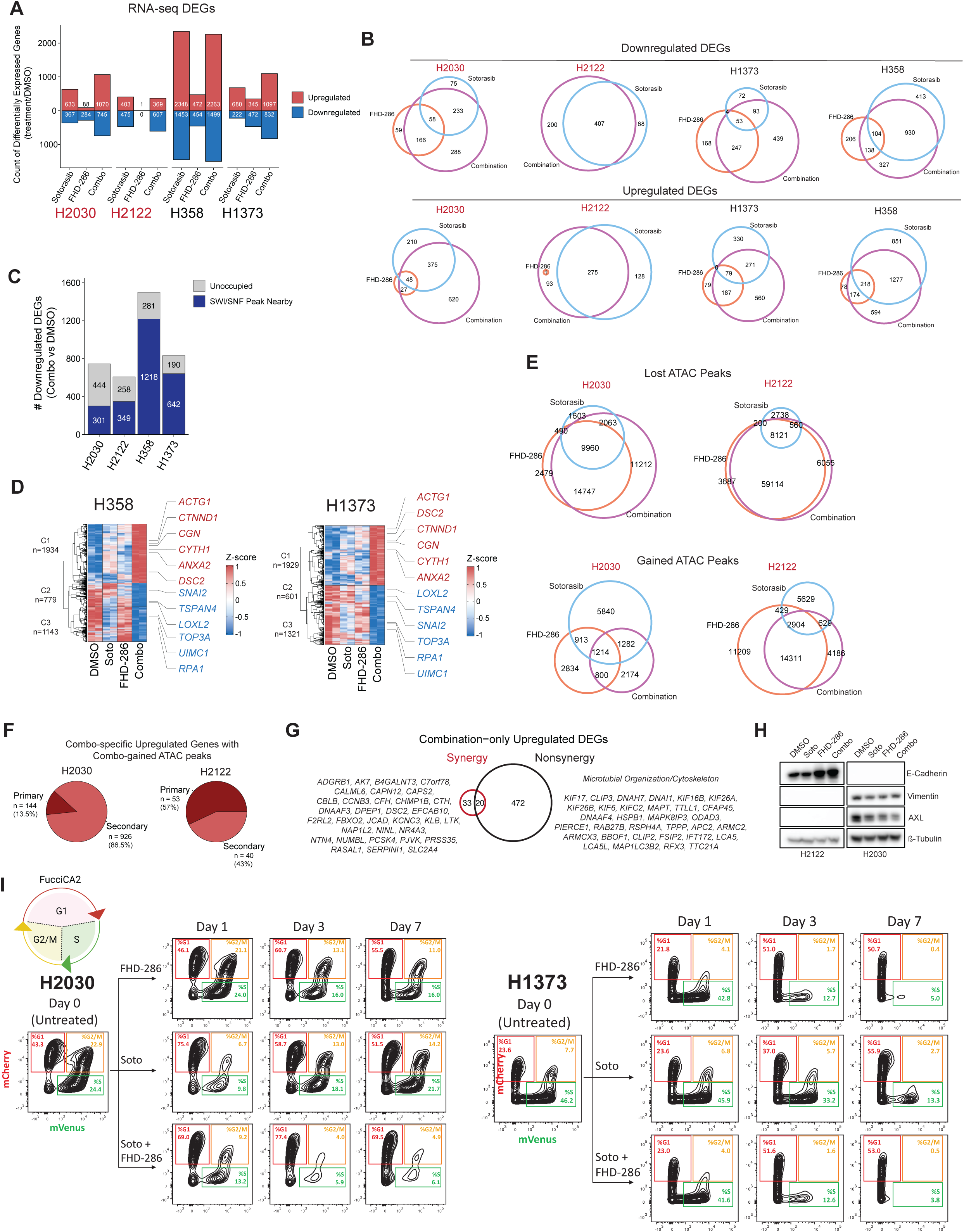
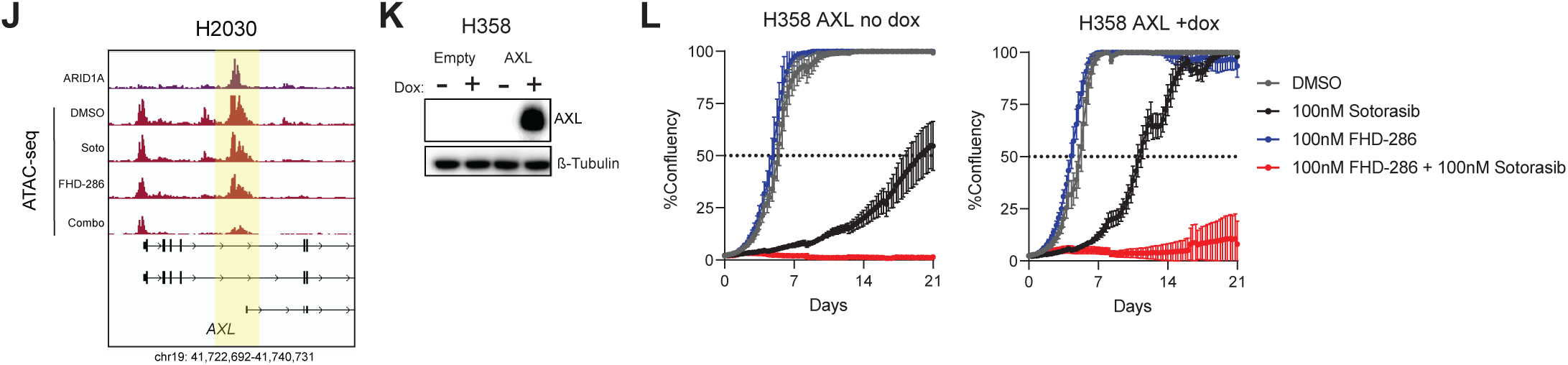
Targeted inhibition of mSWI/SNF complexes in combination with sotorasib treatment impacts EMT and cell cycle across KRAS-mutant cell lines. **A.** Barplots summarizing the number of differentially expressed genes in single and combination drug treatments in synergy and non-synergy cell lines. **B.** Euler diagrams summarizing the number of upregulated or downregulated genes shared between sotorasib, FHD-286 and combination sotorasib and FHD-286 treatments in each cell line. **C.** Barplots summarizing the proportions (with count summaries) of downregulated genes in the combination treatment with mSWI/SNF occupancy in each cell line. **D.** Heatmaps of top 10% deregulated genes upon combination treatment in H358 and H1373 cells. Gene expressoion values were row z-scored normalized. Genes of interest are highlighted in red (up-regulated) or blue (down-regulated). **E.** Euler diagrams summarizing the number of shared and specific lost or gained ATAC-seq chromatin accessible sites following sotorasib, FHD-286 and combination Sotorasib and FHD-286 treatments in each cell line. **F.** Pie charts representing percentage of combination specific upregulated genes with concordant chromatin accessibility (ATAC-seq) changes upon combination treatment (primary targets) in H2030 and H2122. Nearest ATAC-seq peak was assessed within a 30kb window of TSS. **G.** Venn diagram overlap of combination only up-regulated genes in synergy and non-syngery cell lines (resulting from subtracting sotorasib up-regulated genes). Relevant genes are listed. **H.** Immunoblot of EMT markers E-Cadherin, vimentin, and AXL upon sotorasib, FHD-286 or combination drug treatments in H2030 and H2122 cells. B-tubulin is the loading control. **I.** Florescent Ubiquitination-based Cell Cycle Indicator (FUCCI) reporter system in H2030 and H1373 cells following respective treatments at Day 1, 3 and 7. **J.** Representative tracks at the AXL locus in H2030 cells, showing mSWI/SNF complex occupancy (ARID1A CUT&RUN) and chromatin accessibility (ATAC-seq) across treatment conditions. Loss of ATAC signal is highlighted in a yellow box. **K.** Immunoblot of AXL protein levels in H358 cells following 21 days of dox inducible AXL expression. B-tubulin is loading control. **L.** Cell proliferation curves of H358 cells without AXL overexpression (no dox) and with AXL overexpression (+dox) treated with DMSO, sotorasib, FHD-286 or combination following 21 days of treatments.

**Figure S3.**
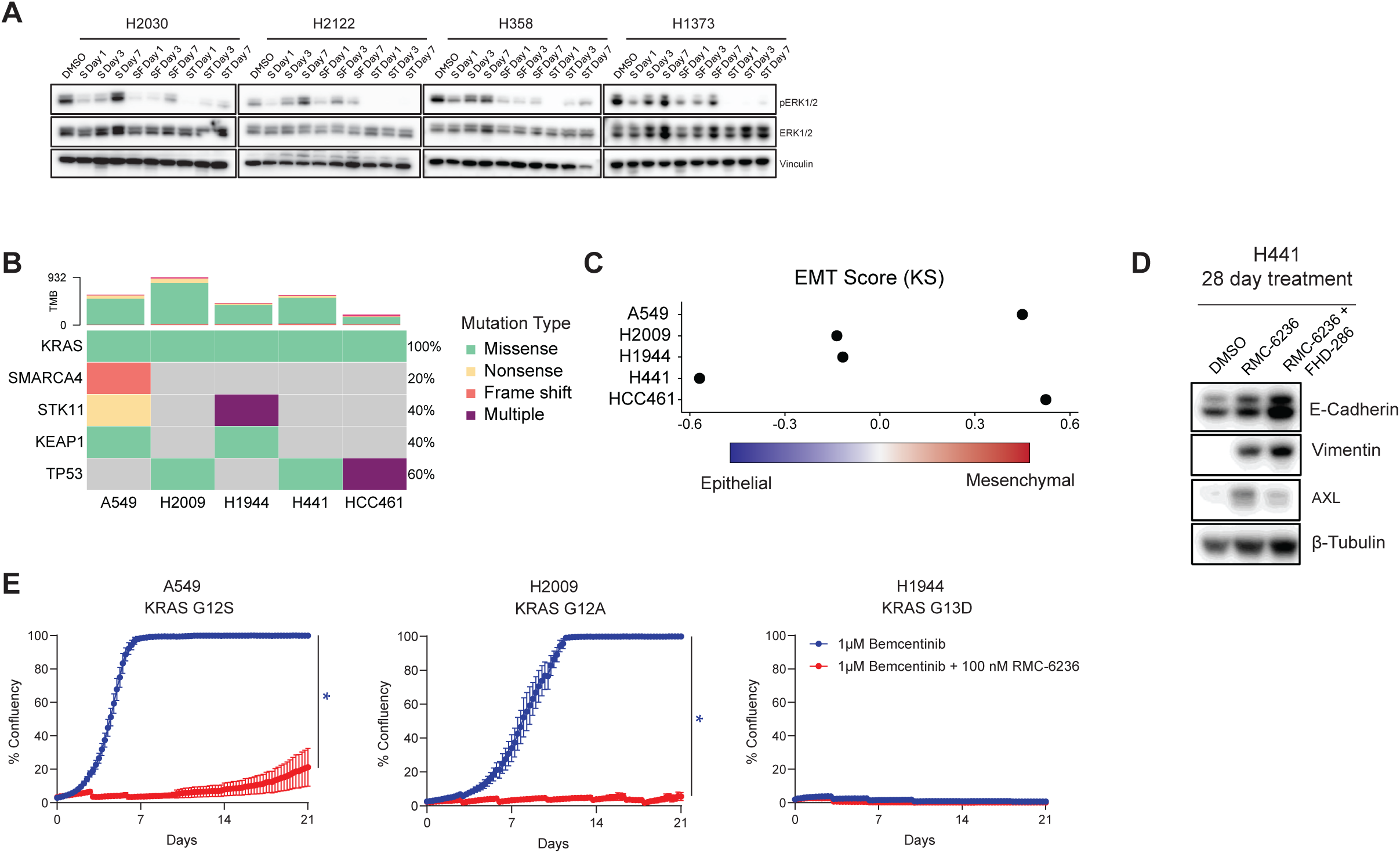
FHD-286 sensitizes non-G12C KRAS-mutant lung cancer models to targeted therapies. A.Western blot of cells treated with DMSO, 100 nM sotorasib (S), 100 nM sotorasib + 100 nM FHD-286 (SF), or 100 nM sotorasib + 30 nM trametinib (MEK inhibitor) (ST) at 1, 3 and 7 days. B. Mutation status of SMARCA4, STK11, KEAP1, and TP53 in non-G12C KRAS-mutant cell lines. C. Epithelial to mesenchymal transition (EMT) score for non-G12C KRAS-mutant cell lines using the KS (Kolmogorov-Smirnov) method. D. Immunoblot of H441 cells treated with 100 nM RMC-6236 for 28 days leading to induction of AXL, which can be blunted by co-treatment with 100 nM FHD-286. E. Confluence of cells exposed to the indicated treatments was measured once every 6 hours over 21 days. Media was replenished every 4 days. Data represents mean +/-SEM of n=4 replicates. Welch’s t-test was used to compare groups at endpoint. *p < 0.05.

**Figure S4.**
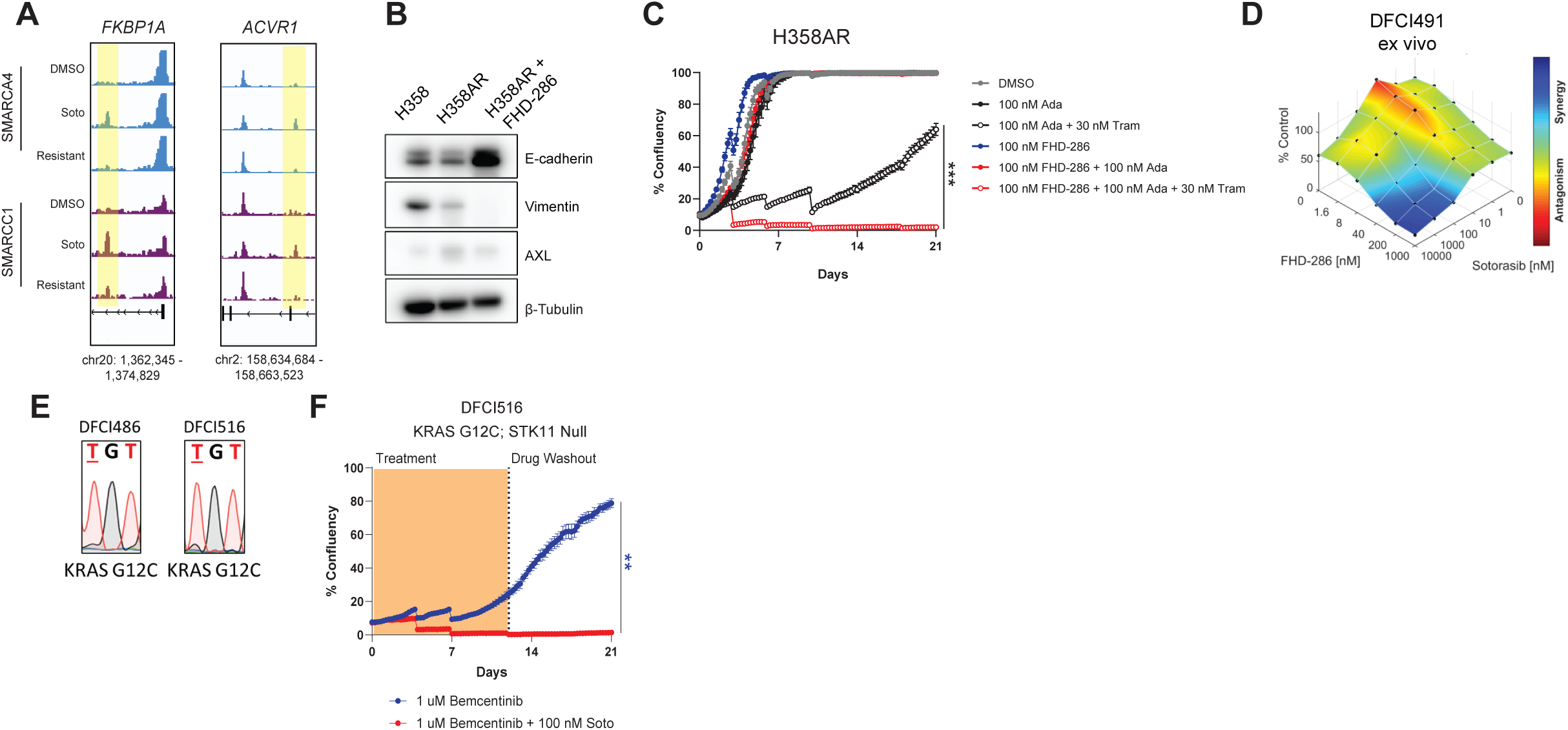
mSWI/SNF inhibition re-sensitizes patient-derived organoid models to KRAS inhibitors. A.IGV track examples of SMARCA4 and SMARCC1 CUT&RUN peaks across DMSO, soto (24hr) and resistant conditions in H358 cells at the *FKBP1A* and *ACVR1* loci. Increased peaks of interest are highlighted. B. Western blot comparing expression of EMT markers in H358, H358AR (in presence of 100 nm adagrasib), and H358AR (in presence of 100 nM adagrasib) treated with 100 nM FHD-286 for 96 hrs. C. Following 1 week adagrasib washout, H358AR cells were challenged with the indicated treatments. Confluence was measured once every 6 hours over 21 days, with media replenished every 4 days. D. DFCI491 ex vivo tumor spheroids were challenged with drug combination matricies with dose titrations of sotorasib and FHD-286 in ULA 384-well plates. Following 96 hours, 3D CTG assay was performed and drug synergy was calculated using Combenefit software. Processed tumor material 40-100 µm was frozen and used for experimental repetitions. One representative experiment out of three is shown. E. Sanger sequencing results for newly established DFCI486 and DFCI516 patient-derived cell lines confirms retention of KRAS G12C mutation. F. Confluence of DFCI516 cells challenged with AXL-specific inhibitor (bemcentinib) alone or in combination with sotorasib. One-way ANOVA was used to compare groups at endpoint. **p < 0.005. Data represent mean +/- SEM of n=4 replicates.

**Figure S5.**
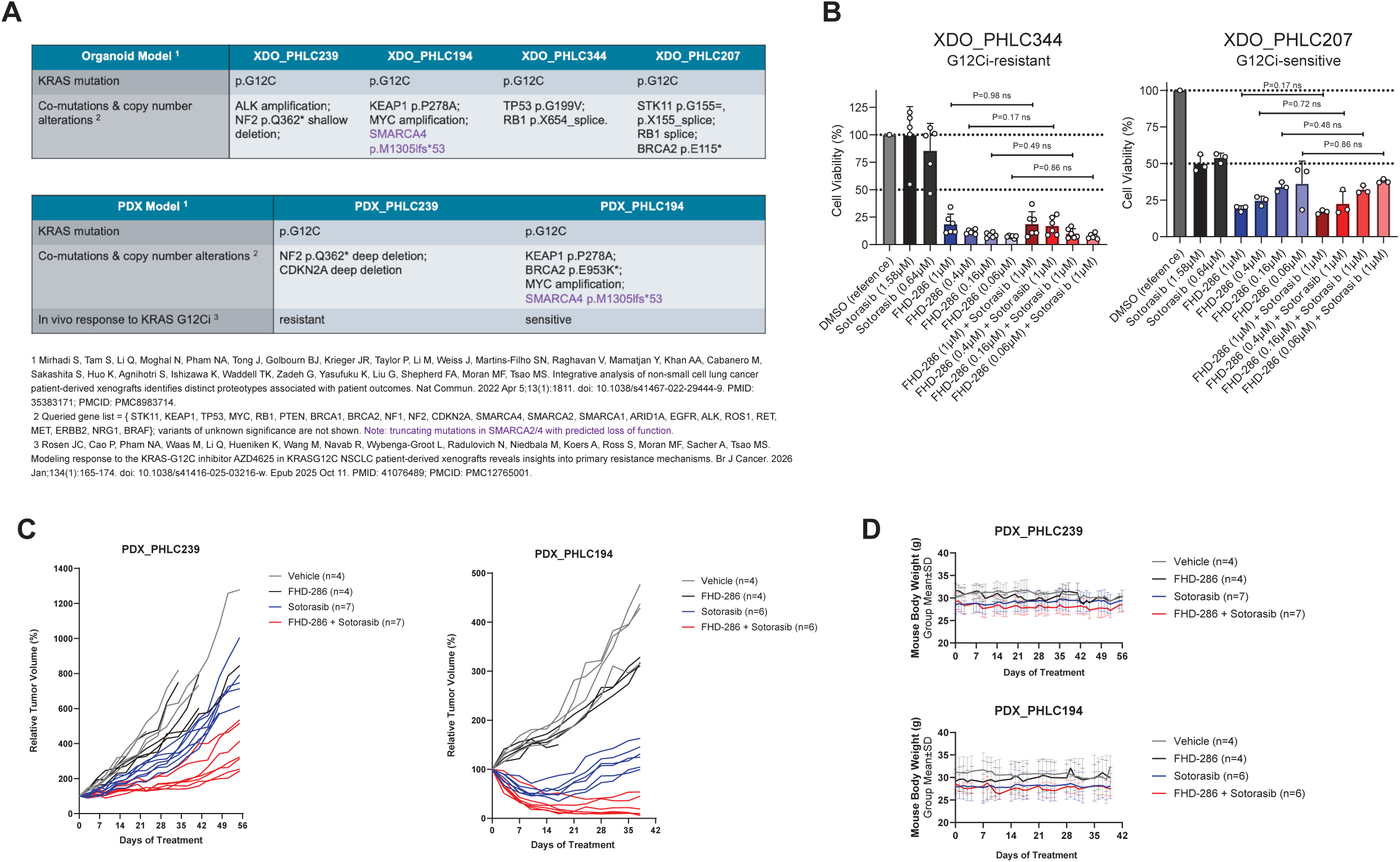
mSWI/SNF inhibition (FHD-286) sensitizes organoid and in vivo patient-derived xenograft models to sotorasib. **A.**Summary table of characteristics for xenograft-derived organoid models (XDOs) and patient-derived xenograft models (PDXs) used in this study. **B.** 3D CTG analysis for organoids, XDO_PHLC344 and XDO_PHLC207, following 14 days of the indicated drug treatments (viability relative to DMSO control). **C.** Individual tumor growth kinetics representative of relative tumor volume for PDX models PHLC239 and PHLC194. **D.** Line graphs of mouse body weight (in grams) across indicated drug treatments in PDX models PHLC239 and PHLC194. Data represent group mean +/- SEM.

